# Bacterial quorum sensing signal arrests phytoplankton cell division and protects against virus-induced mortality

**DOI:** 10.1101/2020.07.14.202937

**Authors:** Scott B. Pollara, Jamie W. Becker, Brook L. Nunn, Rene Boiteau, Daniel Repeta, Miranda C. Mudge, Grayton Downing, Davis Chase, Elizabeth L. Harvey, Kristen E. Whalen

## Abstract

Interactions between phytoplankton and heterotrophic bacteria fundamentally shape marine ecosystems. These interactions are driven by the exchange of compounds, however, linking these chemical signals, their mechanisms of action, and resultant ecological consequences remains a fundamental challenge. The bacterial signal 2-heptyl-4-quinolone (HHQ), induces immediate cellular stasis in the coccolithophore, *Emiliania huxleyi*, however, the mechanism responsible remains unknown. Here, we show that HHQ exposure leads to the accumulation of DNA damage in phytoplankton and prevents its repair. While this effect is reversible, HHQ-exposed phytoplankton are also protected from viral mortality, ascribing a new role of quorum sensing signals in regulating multi-trophic interactions. Further results demonstrate global HHQ production potential and the first *in situ* measurements of HHQ which coincide with areas of enhanced micro- and nanoplankton biomass. Our results support bacterial communication signals as emerging players, providing a new mechanistic framework for how compounds may contribute to structure complex marine microbial communities.

## INTRODUCTION

Interactions between marine phytoplankton and bacteria have been shown to fundamentally shape marine ecosystems, particularly by mediating biogeochemical cycling, regulating productivity, and trophic structure ^1, 2, 3^. Bacteria-phytoplankton interactions are complex, often being species-specific ^4^ or temporally ephemeral ^5^ and can span the spectrum from antagonistic to beneficial ^6, 7^. Increasingly, it is clear that these intricate inter-kingdom interactions are facilitated by excreted chemical compounds that mediate a suite of processes such as nutrient transfer, primary production, and shifts in community composition. Linking chemical compound identity with a mechanism of action and ecological consequences will strengthen our understanding into how these fundamental and multifaceted interactions govern marine ecosystem function.

First discovered in marine systems four decades ago ^8^, quorum sensing (QS) is a form of microbial cell-cell communication through which marine bacteria use diffusible chemical signals to facilitate coordinated and cooperative biogeochemically important behaviors ^9^. Recent work finds alkylquinolone-based QS signals can modulate interspecies behavior, suggesting that these molecules may influence cellular communication at the interkingdom level ^10^. In particular, the alkylquinolone QS signal 2-heptyl-4-quinolone (HHQ) functions as a messenger molecule, able to modulate bacterial virulence behavior, facilitating the emergence of the pathogen *Pseudomonas aeruginosa* within polymicrobial communities ^11, 12^. Additionally, HHQ has also been implicated in antagonizing fungal biofilm formation ^12^, downregulating eukaryotic host immune response via suppression of a key transcription factor, NF-κB ^10^, and activating receptors found to play a role in innate immune signaling in airway epithelia ^13^. These findings support the influence of alkylquinolones in mediating host-microbe interactions. More recently, HHQ was isolated from marine gamma-proteobacteria (*Pseudomonas* sp. and *Pseudoalteromonas* sp.) where it was observed to cause significant shifts in both natural phytoplankton and microbial communities ^14^ and induce species-specific decreases in phytoplankton growth at nanomolar concentrations ^15^. However, the underlying molecular mechanism(s) by which HHQ influences phytoplankton fitness remains unknown.

Here, ultrastructural observations and diagnostic biochemical assays were integrated with transcriptomic and proteomic studies to link the persistent but reversible physiological impact of nanomolar concentrations of HHQ on a model marine phytoplankton, *Emiliania huxleyi* ^15^ in order to determine the molecular underpinnings of HHQ exposure. *E. huxleyi* plays a central role in mediating ocean carbon ^16^ and sulfur ^17^ cycling. Thus, the results presented here emphasize the importance of considering the ecological consequences of chemically-mediated bacteria-phytoplankton interactions for global primary production and biogeochemical cycles.

## RESULTS AND DISCUSION

To detail how HHQ impacts algal growth and morphology, batch cultures of axenic *E. huxleyi* (CCMP 2090) were exposed to 100 ng ml^-1^ of HHQ, a concentration representing the center of a range of inhibitory concentrations (IC_50_) for this species (Fig. S1*A* ^15^). Throughout the remainder of this study HHQ was dosed at a final concentration of 100 ng ml^-1^ unless otherwise specified. Cells exposed to HHQ for 504 h (21 d) exhibited cellular stasis (no cell division nor mortality) concomitant with a significant increase (repeated analysis of variance (ANOVAR), *p*-value < 0.01 for all comparisons) in forward scatter, red fluorescence, and side scatter, proxies for cell size, chlorophyll content, and cell roughness, respectively (Fig. 1*A-C*, Fig. S2*A*). Photosynthetic efficiency (Fv/Fm) did not change in response to long-term HHQ exposure (ANOVAR, Fig. S2*B*). Morphological analysis found *E. huxleyi* cells exposed to HHQ for 24 h had enlarged chloroplasts with distended thylakoid membranes containing numerous intra-organelle vesicles, abundant cytoplasmic vesicles/vacuoles, homogenous nuclei staining lacking defined euchromatin/heterochromatin regions with disintegrated nuclear envelops, and osmium-rich puncta within and adjacent to the chloroplasts likely indicating enhanced lipid storage (Fig. 1*D-E*, Figs. S3, S4). After 14 d of HHQ exposure, cells contained numerous chloroplasts and mitochondria, enhanced cytoplasmic vacuolization, nucleoli with distinct fibrillar centers, abundant lipid droplets, and cultures contained expelled chloroplasts (Fig. 1*F*, Fig. S5). The impact of HHQ on phytoplankton appears to be species-specific, as similar growth and morphology dynamics were not observed for phytoplankton species unaffected by HHQ (Fig. S6; ^15^). Interestingly, when HHQ was diluted to ∼80-fold below the IC_50_ in cultures previously exposed to 100 ng ml^-1^ of HHQ, growth rate, cell size, red fluorescence, and side scatter of *E. huxleyi* recovered to control conditions (ANOVAR, Fig. S7). After 504 h of HHQ exposure, the recovering cells took 144 h to exhibit growth dynamics that were not significantly different to the control, however, recovery did occur, indicating that the effects of HHQ are reversible (ANOVAR, Fig. S7). Recovery or ‘escape’ from growth detrimental conditions has been observed previously for *E. huxleyi* in response to viral infection ^18^.

**Figure 1.**
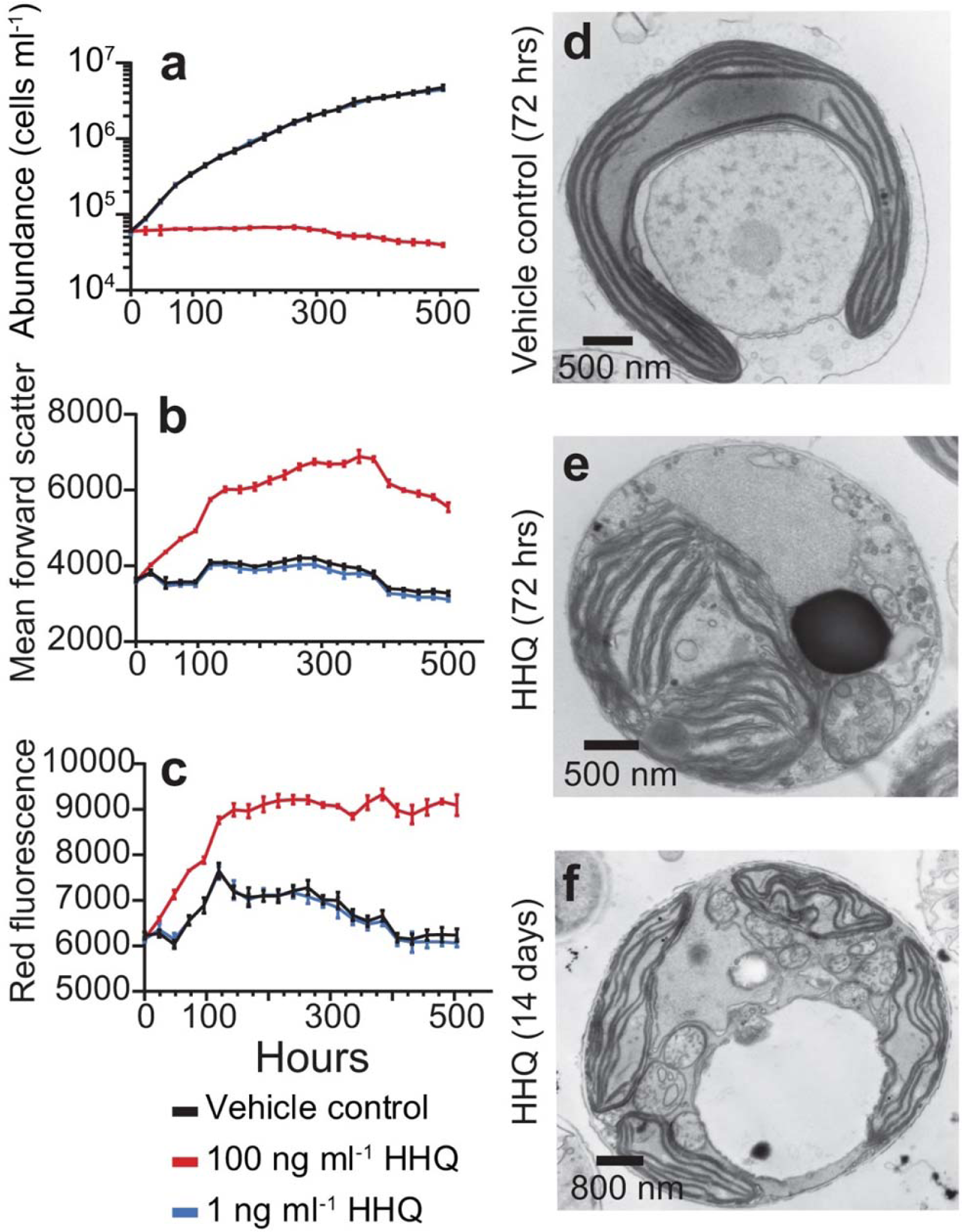
Physiological effects of HHQ on *E. huxleyi. E. huxleyi* cultures (N = 3) were exposed to HHQ or vehicle control (DMSO) and monitored by flow cytometry for cell abundance (a), forward scatter (b), and red fluorescence (c). Mean ± standard deviation shown. Significant differences between HHQ-exposed cells and the vehicle control were evaluated using a repeated measures analysis of variance (ANOVAR; *p* < 0.05). Transmission electron microscopy micrographs of *E. huxleyi* cells exposed to vehicle control (DMSO) (d) or 100 ng ml^-1^ HHQ for 24 h (e), or 100 ng ml^-1^ HHQ for 14 d (f).

To identify eukaryotic pathways affected by HHQ, whole-cell transcriptomic and proteomic analyses were performed on *E. huxleyi* cells exposed to 1 ng ml^-1^ (low), 10 ng ml^-1^ (medium), and 100 ng ml^-1^ (high) HHQ concentrations, with samples taken at 24 h (transcripts) and 72 h (transcripts and proteins). After 72 h of exposure, replicate high HHQ samples appeared distinct from the DMSO vehicle control samples (Figs. S8 – S10), with 37.6% of transcripts (Wald test, *q*-value < 0.05) and 15.9% of proteins (Welch’s approximate t-test, *q*-value < 0.05) significantly changing in relative abundance and abundance, respectively (Figs. S11 and S12, Table S1). When examined together, a total of 665 genes and corresponding proteins were found to be significantly changing in abundance at 72 h under high HHQ treatment relative to the vehicle control (Fig. 2). In general, processes associated with DNA replication and repair, aerobic respiration, and protein catabolism yielded higher relative transcript and protein abundances under high HHQ treatment, while photosynthetic components/processes were detected at lower relative transcript and protein abundances (Fig. 2).

**Figure 2.**
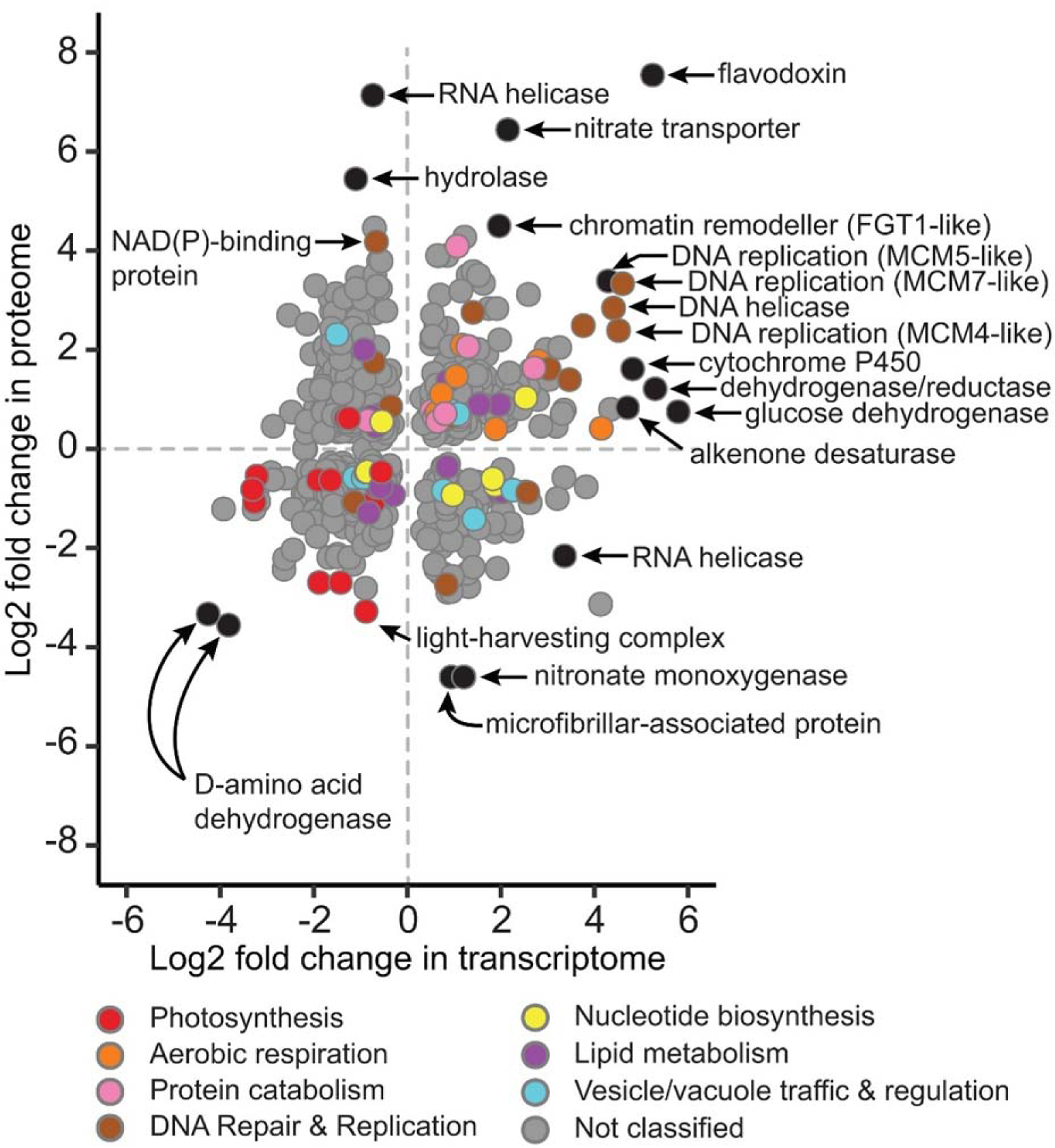
Comparison of log_2_ fold changes in transcript (*x*-axis) and protein (*y-*axis) expression from *E. huxleyi* cultures (N = 4) following exposure to 100 ng ml^-1^ HHQ for 72 h compared to the vehicle control (DMSO). Only shared differentially expressed transcripts (Wald test, *q-*value < 0.05) and proteins (Welch’s approximate t-test, *q*-value < 0.05) are shown. Transcripts and proteins with similar functions are colored via gene ontology (GO) annotation according to the curated groupings shown in Dataset S2. Genes and proteins without GO annotations or annotations outside of the selected groupings are shown in grey. Selected outliers are labeled in black.

### Cell cycle

To investigate the mechanisms of the long-term, but reversible, cellular stasis observed in HHQ-exposed *E. huxleyi*, the impact of HHQ on cell cycle was examined. Using flow cytometry, cell cycle analysis indicated a cessation of the typical cell cycle progression of *E. huxleyi* within 10 h of HHQ exposure, as demonstrated by a gradual accumulation of cells in early S-phase over multiple days (Fig. 3, Fig. S13). The phenotypic response of HHQ treated *E. huxleyi* cells appears to mirror previous studies in which cellular arrest has been observed in phytoplankton in response to bacterially derived chemical exposure ^19, 20, 21, 22, 23^, as well as nutrient limitation ^24, 25, 26^. Indeed, at the physiological level, the response of *E. huxleyi* to HHQ parallels phosphorus (P) limitation in phytoplankton (i.e. S/G2 phase arrest, decreased growth rate, and increases in chlorophyll content, forward scatter, and side scatter) ^24, 25, 27^. However, the canonical response in P-limited cells of upregulation of both alkaline phosphatase and phosphodiesterases ^28, 29, 30^ was not observed in cells exposed to HHQ. Nor do we see significant induction of acid phosphatases, pyrophosphatase, phosphorus transporters, or ATP-sulfurylase enzymes known to be induced following P-limitation in HHQ exposed cells, indicating the lack of phosphorus stress (Dataset S1). Therefore, while the pattern of cell cycle arrest is similar between HHQ-treated *E. huxleyi* and nutrient limitation, the underlying mechanisms are distinct.

**Figure 3.**
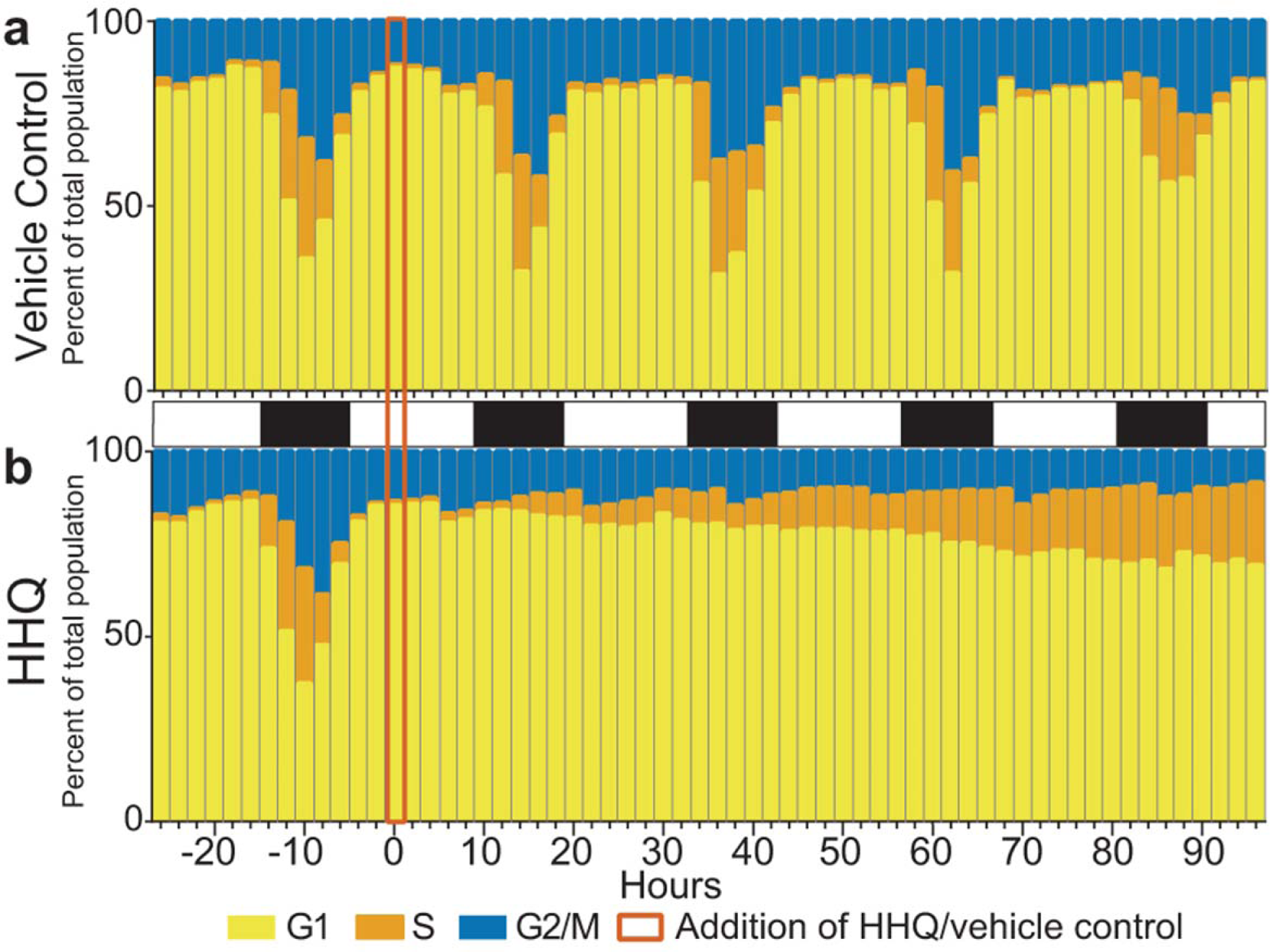
Quantification of cell cycle stage following HHQ exposure. *E. huxleyi* cultures (N = 3) were exposed to either vehicle control (DMSO) (a) or 100 ng ml^-1^ HHQ (b) for 96 h and their DNA content was profiled to determine their cell stage.

In phytoplankton, cellular arrest is often accompanied by induction of autocatalytic or programmed cell death (PCD) responses such as increased reactive oxygen production or caspase-like activity ^31^, and previous findings in mammalian cells indicate that HHQ has the ability to activate PCD pathways ^32^. Using a series of diagnostic fluorescent assays (i.e. membrane permeabilization, caspase activity, reactive oxygen species (ROS), and nitrous oxide (NO) production) (Fig. S14) on HHQ exposed *E. huxleyi* cells, no evidence of PCD/apoptosis was discovered. Additionally, no transcripts or proteins associated with PCD increased in abundance with exposure to HHQ (Dataset S1). The occurrence of cellular arrest in the absence of apoptosis indicates that HHQ exposed cells likely progress through the G1/S transition commitment point, but then stall and accumulate in S-phase (Fig. 3*B*, Fig. S13). This observation is supported by the increased relative abundance of canonical transcripts enabling the G1/S transition including cell division control protein 6 (CDC6), origin recognition complex subunit 1 (ORC), and cyclins A, B, E, and K, in HHQ-exposed treatments (Fig. 4, Dataset S1). Moreover, significant increases in relative transcript abundances of DNA replication fork machinery (i.e., DNA polymerases α, ε, and δ, DNA primase, replication protein A, topoisomerases (TOPO), the minichromosomal maintenance complex, proliferating cell nuclear antigen, and replication factor C, Fig. 4; Dataset S1) at 72 h post HHQ exposure, suggests an intent to replicate DNA, a hallmark of S-phase ^33^. However, despite this observed induction of DNA replication machinery, cell cycle analysis demonstrated DNA synthesis was severely diminished following HHQ exposure (Fig. 3, Fig. S13), suggesting that HHQ interferes with the ability of *E. huxleyi* cells to correctly complete the DNA replication process. Disruption of DNA replication induces DNA damage response pathways thereby activating effector kinases such as Chk1 and Chk2 (Fig. 4) necessary for the halting of DNA synthesis and induction of cell cycle arrest to allow for time for repair ^34^. Under HHQ treatment, Chk1 and Chk2 are differentially abundant, however, protein phosphorylation patterns need to be interrogated to understand how these DDR regulators are impacted by HHQ presence. Moreover, we observed a significant decrease in the relative abundance of histone transcripts and proteins (Fig. 4) following HHQ exposure, which is a hallmark of DNA synthesis disruption as DNA replication and histone production are coupled and the cell possesses pathways to remove histone transcripts following DNA replication stress ^35^.

**Figure 4.**
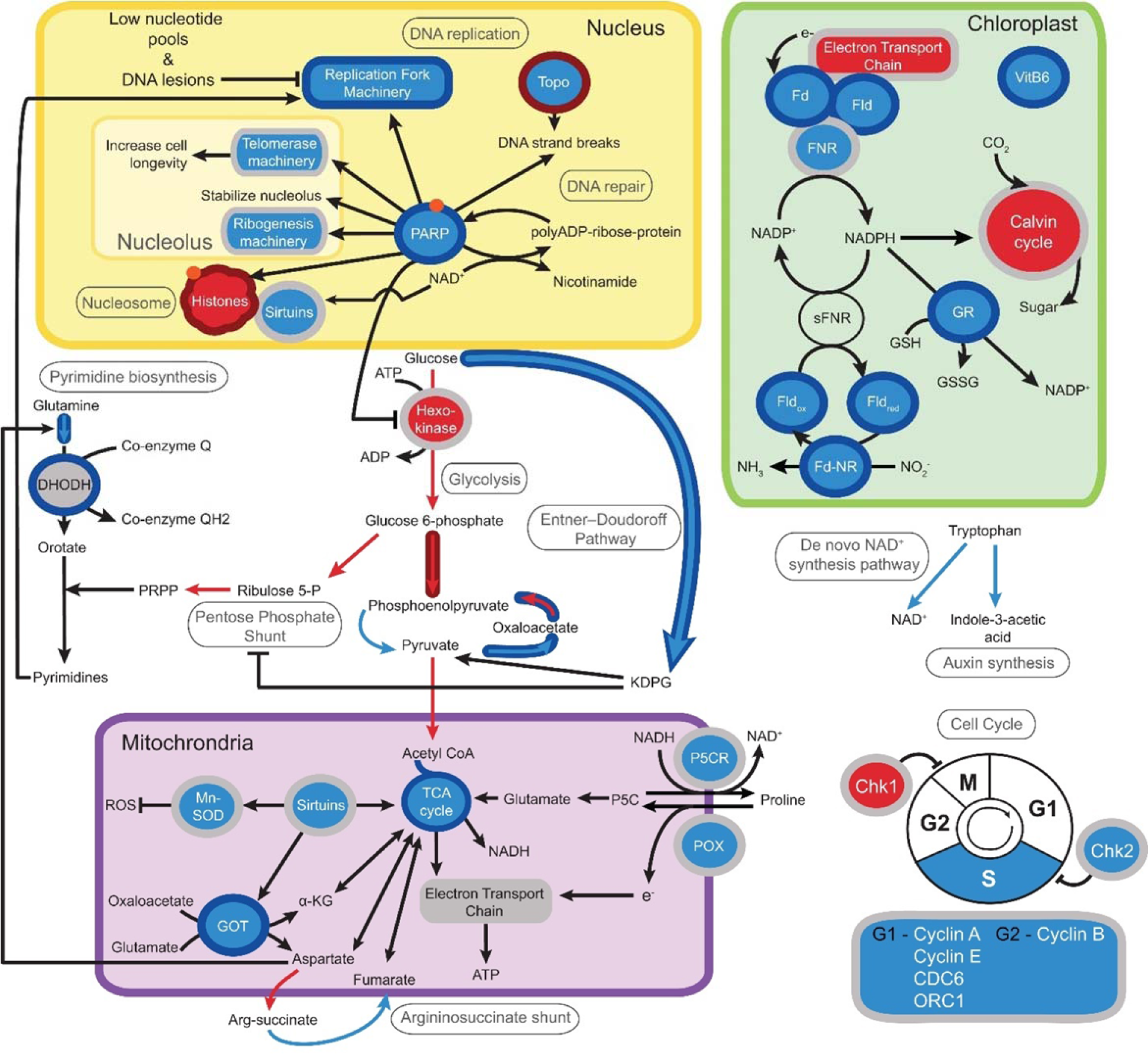
Schematic displaying the significant increases (blue) and decreases (red) in transcript (Wald test, *q-*value > 0.05) and protein (Welch’s t-test, *q*-value < 0.05) abundance following HHQ exposure in *E. huxleyi* cultures relative to vehicle control (DMSO). The interior color of shapes and arrows denote significant transcript changes, while the outline color of shapes or arrows denotes significant protein changes. Grey shapes or outlines indicate no differential expression was noted at any time point. [Nucleus] *PARP*: Poly (ADP-ribose) polymerase; *TOPO*: Topoisomerase. [Chloroplast] *Fd*: Ferredoxin; *FNR*: Ferredoxin-NADP+ oxidoreductase; *Fd-NR*: Ferredoxin nitrite reductase; *VitB6*: pyridoxine biosynthesis protein; *sFNR*: soluble ferredoxin-NADP+ oxidoreductase; *Fld*: Flavodoxin, *GR*: Glutathione reductase; *GSH*: Glutathione. [Cytosol] KDPG: 2-Keto-3-deoxy-6-phosphogluconate; PRPP: Phosphoribosyl pyrophosphate; *DHODH*: Dihydroorotate dehydrogenase. [Mitochondria] TCA cycle: Tricarboxylic acid cycle; *Mn-SOD*: Manganese superoxide dismutase; ROS: Reactive oxygen species; *GOT*: Glutamic oxaloacetic transaminase; α-KG: Alpha-ketoglutarate; *P5C*: pyrroline-5-carboxylate; *P5CR*: pyrroline-5-carboxylate reductase; *POX*: Proline oxidase. [Cell Cycle] *Chk1*: Serine/threonine-protein kinase Chk1; *Chk2*: Serine/threonine-protein kinase Chk2; *CDC6*: Cell division control protein 6; *ORC1*: Origin recognition complex subunit 1. Orange dots indicate potential targets of parylation by PARP proteins.

### DNA replication and repair

During S-phase, a cell must tightly regulate the availability of nucleotides to ensure faithful DNA replication ^36^. Therefore S-phase cells rely on *de novo* nucleotide synthesis pathways to produce enough materials for complete genome replication ^37^. Several transcripts and proteins involved in *de novo* purine (amidophosphoribosyltransferase, trifunctional purine biosynthetic protein adenosine-3 (GART), and phosphoribosylformylglycinamidine synthase, bifunctional purine biosynthesis protein (ATIC), adenylsuccinate synthase, IMP dehydrogenase, and GMP synthase) and pyrimidine (carbamoyl phosphate synthase II, aspartatecarbamoyl transferase, and CTP synthases) nucleotide synthesis increased in abundance with HHQ exposure (Fig. 4, Dataset S1). Increased nucleotide synthesis may indicate the need to produce the necessary materials to replenish nucleotide pools during replication. However, we observe only partial replication of the *E. huxleyi* genome following HHQ exposure (Fig. 3, Fig. S13), suggesting nucleotide availability is limited and HHQ may disrupt the production of nucleotides.

Alkylquinolones are known to inhibit a key rate-limiting enzyme directly involved in bacterial pyrimidine synthesis, dihydroorotate dehydrogenase (DHODH) ^38^. DHODH inhibition in eukaryotes may induce an intra-S-phase arrest due to severely diminished cellular nucleotide pools that can disrupt DNA replication, stall replication forks, and increase the frequency of genomic DNA lesions, including strand breaks, during S-phase ^39, 40^. Indeed, after 46 h of HHQ exposure, a significant increase in DNA strand breaks was observed in culture (Welch’s approximate t-test, *p* = 0.032; Fig. S15*A*), and not observed when HHQ was directly exposed to genomic *E. huxleyi* or Lambda DNA (Fig. S15*B*). This indicates that DNA strand breaks are not caused directly by HHQ, but indirectly through other mechanisms. It has been previously observed that following the induction of DNA damage during S-phase, cells will enter an intra-S phase arrest that drastically slows the rate of DNA replication to allow the DNA damage response (DDR) to resolve any DNA lesions ^41^. With the exception of preliminary work in *Chlamydomonas reinhardtii* and dinoflagellates, the DDR response has not been well characterized in phytoplankton ^42, 43^. Of the 57 mammalian DDR protein homologs in the *E. huxleyi* genome (e-value ≤ 10^−20^), 41 were significantly differentially expressed, of which 37 increased in relative abundance at 72 h under high HHQ exposure (Dataset S1), indicating the cell is attempting to repair DNA lesions. However, DNA damage induced by the inhibition of DHODH is known to activate apoptotic pathways through the hyperactivation of the DDR ^44^. No apoptotic pathway activation was observed with HHQ exposure, suggesting the DDR response itself may also be impacted by HHQ.

A master regulator of the DDR involved in chromatin remodeling, nucleolar structure (thereby facilitating formation of telomerase and ribogenesis machinery), and genome stability is poly(ADP-ribose) polymerase (PARP) (Fig. 4) ^45^. PARP binds to sites of DNA damage and stalls replication forks and produces negatively charged ADP-ribose polymers to serve as a scaffold for the necessary repair proteins to resolve the DNA lesion or restart the fork ^46^. PARP homologs in *E. huxleyi* were found to increase in both relative transcript abundance and protein abundance under HHQ treatment (Dataset S1). Further, HHQ alteration of *E. huxleyi* nucleoli morphology (Figs. S4, S5) and the differential abundance of genes and proteins in PARP activity-dependent processes, including ribogenesis and telomerase biogenesis (Dataset S1; ^47, 48^), was observed. Interestingly, the genomes of phytoplankton species unaffected by HHQ ^15^ did not reveal the presence of any PARP homologs, further implicating PARPs in the response of phytoplankton to HHQ.

Under high levels of DNA damage or if repair mechanisms are compromised, PARP can become overactivated and deplete cellular NAD^+^ and ATP pools, thereby inducing apoptotic pathways ^49^. As no apoptotic activity was observed in these experiments, HHQ may inhibit PARP activity. Inhibition of PARP activity in the presence of DNA damage drastically reduces the effectiveness of the DDR response and is known to induce cellular arrest in the S-phase ^50^. HHQ was found to significantly inhibit human PARP activity (Welch’s approximate t-test, *p* = 0.0002; Fig. S15*C*), while a closely related alkylquinolone, 2-heptyl-3-hydroxy-4(1H)-quinolone (PQS), did not possess PARP inhibitory activity, nor did it impact *E. huxleyi* growth (Figs. S1*B* and S15*B*). Together, the observation of prolonged S-phase arrest, the upregulation of the DDR response in HHQ-exposed cultures, the conserved nature of the mammalian and *E. huxleyi* PARP catalytic site (Fig. S16), and the chemical structural similarities of HHQ to known inhibitors of both PARP and DHODH with core benzimidazole moieties ^51^, collectively suggest that HHQ may function simultaneously to inhibit both PARP and DHODH activity in *E. huxleyi*. Additional experiments using *E. huxleyi* enzymes are needed to fully characterize whether PARP and DHODH are molecular targets of HHQ.

### Energy Production

In order to facilitate DNA synthesis and repair, the cell requires large ATP pools ^52^, and there were several lines of evidence to support ATP generation in HHQ exposed cells. First, an increased relative transcript abundance of enzymes in the tricarboxylic acid (TCA) cycle (i.e., isocitrate dehydrogenase, α-ketoglutarate dehydrogenase, succinate dehydrogenase, fumarase, and malate dehydrogenase) (Fig. 4, Dataset S1) was observed in HHQ exposed cells, indicating the potential overproduction of reducing equivalents for ATP production via oxidative phosphorylation. Second, HHQ exposure had a profound effect on ATP/ADP transport, as transcripts for an ATP/ADP translocase, which catalyzes the highly specific transport of ATP across membranes in exchange for ADP, increased in relative abundance (Dataset S1). Third, an increase in the relative transcript abundance of sirtuin-like deacetylases, metabolic efficiency controllers ^53^, were observed following HHQ exposure (Fig. 4, Dataset S1). Sirtuins compete with PARPs to use NAD^+^ and expression of these deacetylases is dependent on NAD^+^ availability ^54^. PARP inhibition is known to drastically increase cellular NAD^+^ pools, thereby promoting sirtuin expression and activity ^55^. Increased sirtuin activity in HHQ exposed cells may also explain the increase in the relative transcript abundance of manganese superoxide dismutase (Mn-SOD) (Fig. 4), an antioxidant enzyme that protects the cell from ROS induced damage ^56^. Finally, increased relative transcript abundance of the tryptophan-mediated *de novo* NAD^+^ synthesis pathway was also observed, potentially in an attempt to increase NAD^+^ availability (Fig. 4, Dataset S1). Taken together, these results suggest that HHQ exposure promotes increased energy production in *E. huxleyi*, which may enable the cell to fuel various biosynthesis and repair pathways while staving off the induction of PCD.

### Aspartate metabolism

In HHQ exposed cells, an increase in relative transcript abundance was observed for multiple pathways leading to the production of aspartate (i.e., TCA cycle, the aspartate-arginosuccinate shunt, glutamic oxaloacetic transaminase (GOT), and C4-like photosynthesis; Fig. 4, Dataset S1). In parallel, a decrease in the relative abundance of transcripts for aspartate utilization pathways, with the exception of nucleotide synthesis pathways, was observed (Fig. 4, Dataset S1). Aspartate is known to rescue cells from S-phase arrest by fueling *de novo* nucleotide synthesis ^57^. Another way the cell can recover amino acids, including aspartate, is through the degradation of proteins via proteasomal pathways ^58^, which increased on both the transcript and protein level following HHQ treatment (Fig. 2). While the increased cellular demand for ATP would necessitate the upregulation of glycolytic enzymes like hexokinase, the first step in glycolysis, there was a significant decrease in the relative transcript abundance of hexokinase (Fig. 4). These findings are consistent with previous work demonstrating alkylquinolones suppress induction of this glycolytic enzyme through the direct targeting of the transcription factor hypoxia-inducible factor 1 (HIF-1) protein degradation via proteasomal pathways ^59^. Furthermore, in order to conserve amino acid resources, a shift to the Entner-Doudoroff glycolytic pathway in HHQ treated cells (Fig. 4, Dataset S1) was observed. The Entner-Doudoroff glycolytic pathway has a lower protein demand in comparison to other glycolytic pathways ^60^. These results suggest that *E. huxleyi* may be shunting resources from various pools towards the production of aspartate to alleviate the effects of HHQ induced S-phase arrest. However, further investigations measuring the availability of aspartate and aspartate-derived metabolites in HHQ exposed cells is required to understand what is being produced following this shift in *E. huxleyi* metabolism.

### Photosynthesis and redox

HHQ-induced cell cycle arrest in *E. huxleyi* did not significantly alter photosynthetic energy conversion efficiency, however, the majority of light-harvesting complexes and transcripts of the Calvin cycle decreased in relative abundance under HHQ exposure (Figs. 2 and 4). These findings parallel those described for the diatom *Phaeodactylum tricornutum* undergoing chemically-mediated cell cycle arrest ^61^. In plants, the coordinated down-regulation of transcripts involved in photosynthesis, electron transport (i.e. photosystem I and II, ATP synthase, and light-harvesting complexes), and the Calvin cycle is thought to allow for the reallocation of resources towards defense against bacterial and viral pathogens ^62^. However, a decrease in transcript abundance does not always correlate with a loss of function. Photosynthetic proteins have a long functional half-life in the cell with the exception of ferredoxin (Fd) and ferredoxin NADP+ oxidoreductase (FNR) ^62^. Following pathogen infection, cellular redox state can be altered eliciting an increase in both transcript abundance and protein expression of ferredoxin and FNR ^62^. The maintenance of cellular redox pools for metabolism and antioxidant defense requires an influx of electrons via light-based reactions and NADPH-powered redox cascades. In photoautotrophs, all reducing power derived from photosynthetic electron transport passes through ferredoxin acting as an electron distribution hub able to provide feedback on the redox state of the chloroplast ^63^. Together, both ferredoxin and the isofunctional flavodoxin (Fld) participate in electron shuttling between cellular sources of reducing power and electron-consuming routes, preventing electron misrouting that can lead to ROS accumulation and restoring chloroplast redox homeostasis under environmental stress ^64^. Indeed, the genes and proteins with the most significant differential expression levels under HHQ exposure in *E. huxleyi* were Fd (58-fold increase in transcript and 3-fold increase in protein), FNR (85-fold increase in transcript), and Fld (38-fold increase in transcript and 186-fold increase in protein) (Fig. 2 & 4, Dataset S1), which may explain the observed lack of ROS production (Fig. S14*C-E*). Additional reduction systems including FAD/NAD(P) oxidoreductases, ferredoxin nitrite reductase (Fd-NR), and glutathione reductase (GR) in HHQ treated *E. huxleyi* were also significantly induced which could ameliorate NADPH build-up (Fig. 4). Moreover, HHQ exposure resulted in an increased relative expression of vitamin B6 (VitB6) transcripts, which has been shown to protect against oxidative stress in chloroplasts (Fig. 4) ^65^. In addition, under HHQ treatment transcripts for proline oxidase (POX) and pyrroline-5-carboxylate reductase (P5CR) increased in relative abundance (Fig. 4). POX is involved in protection against metabolic stress, and is closely linked with the TCA cycle by donating electrons to the electron transport chain to support ATP generation. Furthermore, both POX and P5CR contribute to the cycling of proline between the cytosol and mitochondria eventually leading to ATP generation and NAD^+^ production to maintain redox homeostasis ^66^ or to be used by other enzymes, possibly PARP. Together, these results suggest that HHQ exposed *E. huxleyi* uniformly decreased the relative abundance of photosynthetic gene transcripts in support of a coordinated induction of defense responses aimed at maintaining cellular redox homeostasis without debilitating photosynthetic capacity.

### Auxin Production

The small signaling molecule indole-3-acetic acid (IAA) is an abundant plant hormone known to control plant and algal growth and cell division ^67^. Homologs of the genes involved in several tryptophan-dependent IAA biosynthesis pathways were found to increase in relative abundance under HHQ treatment (Fig. 4). Previous studies have indicated only those coccolith-bearing *E. huxleyi* strains were capable of producing IAA, however, naked strains, like those used in this study, were more susceptible to IAA effects including increasing cell size and impaired membrane integrity ^67^. Interestingly, IAA at concentrations up to 25 µg ml^-1^ significantly stimulated biofilm formation in the HHQ-producing bacteria, *Pseudomonas aeruginosa* (PAO1) and suppressed growth of planktonic bacterial cells ^68^. Additional work is necessary to investigate if IAA produced by *E. huxleyi* acts as a bacterial attractant, thereby invoking similar biofilm behaviors in associated *Pseudoaltermonas* strains.

### Ecological Consequences

Given that viral replication requires hijacking of host-replication machinery and HHQ exposure inhibited DNA replication in *E. huxleyi*, the impact of HHQ on host-virus dynamics was investigated. When *E. huxleyi* cells were exposed to HHQ and *E. huxleyi* virus (EhV) strain 207, virus-induced cellular death was significantly reduced (ANOVAR, *p-*value < 0.0001; Fig. 5). Visual inspection of electron microscopy images showed enhanced lipid content and cytoplasmic vacuoles/vesicles following HHQ treatment (Fig. 1*E-F*, Figs. S4, S5) in parallel with transcriptional induction of cellular components that mediate endocytosis (i.e., alpha-adaptin, dynamin), vacuolar/vesicle trafficking (i.e., vesicle protein sorting; VPS45, vacuolar sorting protein 46A, coatomer protein, ADP ribosylation factor), membrane traffic (i.e., Rab GTPase), and cytoskeletal components/modifiers (i.e., tubulin, cofilin) including motors (i.e., kinesin) (Dataset S1). This suggests that HHQ may impact the entry and delivery of eukaryotic viral DNA to a host cell. Viral infection will significantly alter the metabolism of the infected cell and influence the production of metabolites, altering the landscape of metabolites available for uptake ^69^. Protection against viral mortality, would theoretically permit increased survival of phytoplankton and allow for bacteria to continue to take advantage of coordinated nutrient exchange, common between bacteria and phytoplankton ^70^. To better understand these ecological consequences, additional work is needed to clarify whether HHQ-induced protection against viral mortality is due to a decrease in infection, a cessation of viral replication, or the inability of the viral particles to lyse the cell.

**Figure 5.**
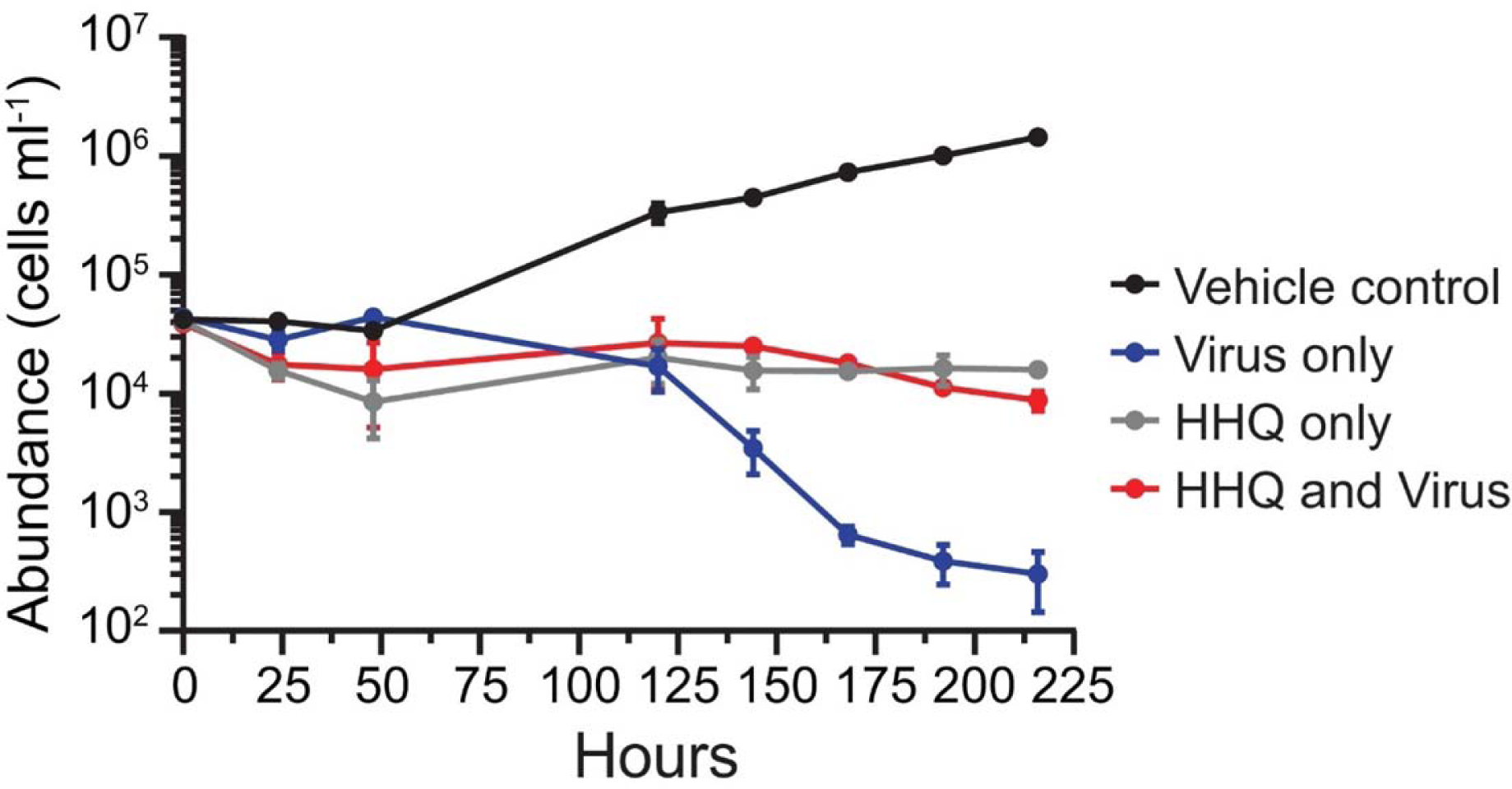
Inhibition of viral-induced mortality in the presence of HHQ. *E. huxleyi* cultures (N=3) were exposed to either vehicle control (DMSO) or 100 ng ml^-1^ HHQ in the presence and absence of viruses. Phytoplankton cellular abundance was measured via flow cytometry and evaluated using ANOVAR (*p* < 0.05). Mean ± standard deviation shown.

### Broader significance and conclusions

Finally, the distribution of HHQ in surface waters and prevalence of HHQ biosynthetic machinery was examined to understand the potential ecosystem level impact of this bacterial QS molecule. Using the TARA Ocean Gene Atlas web service, the signatures of eight genes involved in alkylquinolone synthesis were found to be globally distributed - extending from surface waters to the deep chlorophyll maximum layer (Fig. S17), indicating the potential for HHQ synthesis is ubiquitous. Additionally, LC-ESI-MS chemical analysis found > 1 ng L^-1^ surface concentrations of HHQ in the eastern tropical South Pacific, while concentrations were below the limit of detection (< 0.18 ng L^-1^) in open ocean oligotrophic waters (Fig. 6; Fig. S18). Although these measured bulk concentrations were well below the IC_50_ for coccolithophores, they likely do not represent the effective concentration a marine microbial cell would experience in the phycosphere ^71^. Evidence from the biomedical literature indicates alkylquinolones, including HHQ, can be concentrated to micromolar levels within membrane vesicles and directly released by bacteria ^72^. For marine bacteria closely associated with phytoplankton, these membrane vesicles could directly expose phytoplankton cells to concentrations of HHQ well beyond the experimental concentrations used here. While environmental genomic data and bulk chemical analysis provide important insights into the ubiquity of potential phytoplankton-bacterial interactions, quantifying bacterial metabolites in the phycosphere microenvironment remains a critical challenge to understanding the role of secondary metabolites in phytoplankton-bacterial interactions ^71^.

**Figure 6.**
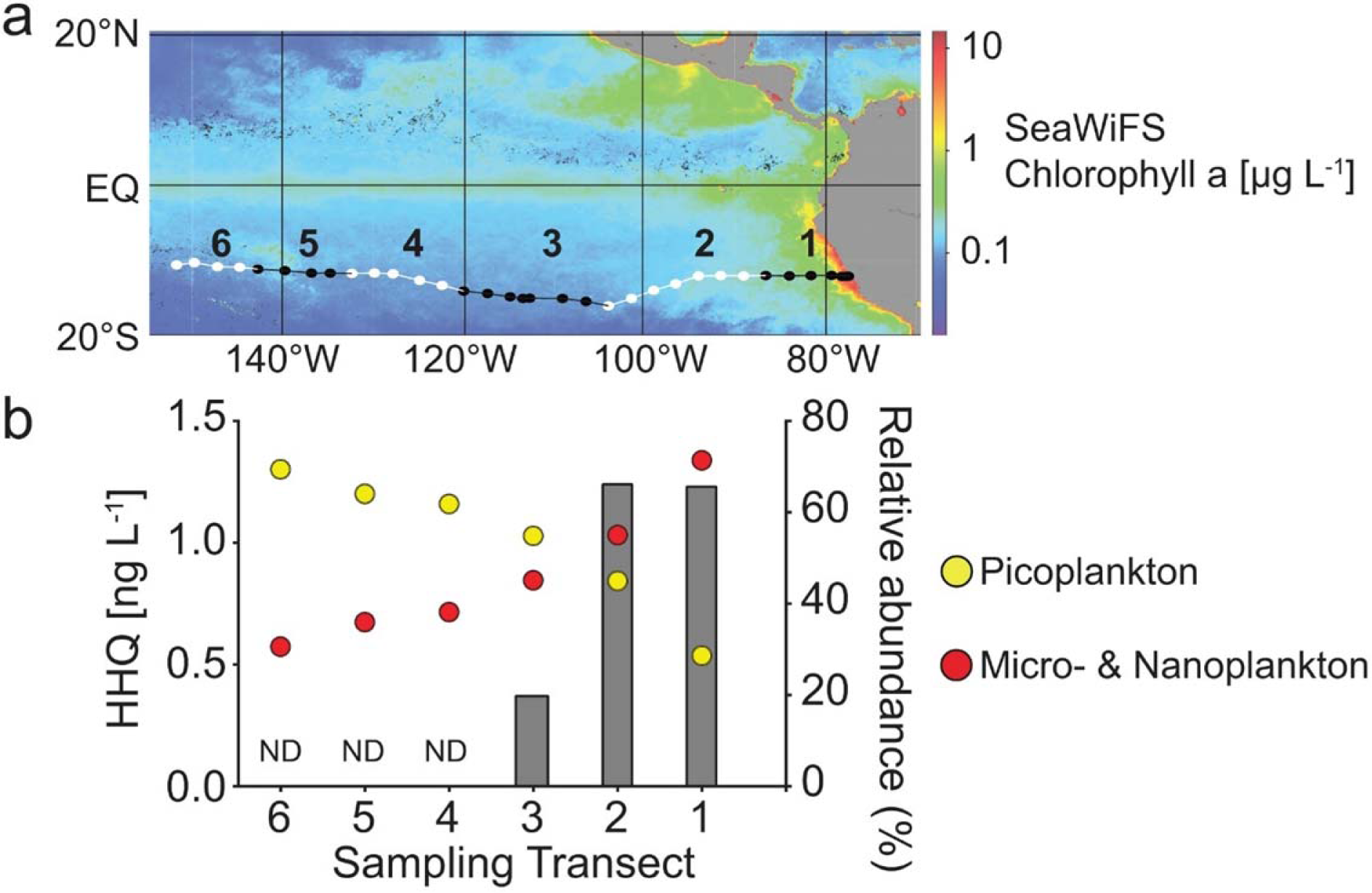
Detection of HHQ in the marine environment. (a) Cruise track of the U.S. GEOTRACES GP16 cruise in 2013 in the eastern southern tropical Pacific Ocean. (b) Grey bars indicated the concentration of HHQ from six stations along the cruise track. Circles indicate the percent relative abundance picoplankton (yellow) and micro- & nanoplankton (red).

These findings demonstrate that a quorum sensing signal produced by a marine bacterium significantly, but reversibly, leads to DNA lesions in a eukaryotic phytoplankter, thereby arresting the cell cycle, reprograming intermediate metabolism, and restructuring cellular architecture therefore significantly influencing inter-kingdom interactions in the sea. As a consequence, these chemically-induced changes are demonstrated to have a cascading impact on a major vector of phytoplankton death, viral mortality. Given the ubiquity of HHQ synthesis genes and relative abundance of HHQ in the marine environment coinciding with enhanced phytoplankton biomass, our results suggest that alkylquinolone signaling plays a significant role in structuring complex microbial communities and, ultimately, influencing primary production and biogeochemical cycles. In addition, our findings highlight the functional duality of bacterial cues that serve both as diffusive messengers used as a communication tool in microbial communities, but also as chemical mediators of eukaryotic physiology capable of impacting trophic level dynamics in marine ecosystems.

## MATERIALS AND METHODS

### General Cultivation Conditions

Three species of phytoplankton were used in this study, axenic *Emiliania huxleyi* (CCMP2090), *Phaeodactylum tricornutum* (CCMP2561), and *Dunaliella tertiolecta* (CCMP1320); all from the National Center for Marine Algae and Microbiota, East Boothbay, Maine) were grown in natural seawater-based f/2 medium both with (*P. tricornutum*) and without (*E. huxleyi* and *D. tertiolecta*) silica ^73^. Cultures were maintained on a 14:10 h light (80 ± 5 µmol photons m^-2^ s^-1^):dark cycle at 17 °C, and salinity of 35. These conditions will be referred to hereafter as general culturing conditions. Strain purity was confirmed using f/2 MM and f/2 MB purity test broths (Table S2) and visually confirmed by epifluorescence microscopy ^74^. Cultures were transferred weekly to maintain exponentially growing cultures.

Phytoplankton cells were enumerated by hemocytometer or using a flow cytometer (Guava, Millipore). Via the flow cytometer, cell abundance was determined by using species-specific settings including their forward scatter, side scatter, and red fluorescence (695/50 nm) emission characteristics. All samples were run at 0.24 µl s^-1^ for 3 min, either live or fixed with glutaraldehyde (0.5% final concentration). A correction factor was applied to fixed cell abundances to account for cell loss due to preservation.

### Growth Experiments

The HHQ concentration resulting in 50% growth inhibition (IC_50_) was determined using triplicate 2 - 20 ml cultures of *E. huxleyi* (∼100,000 cells ml^-1^) exposed to HHQ (between 0.25 – 512 ng ml^-1^), PQS (0.5 – 530 μg ml^-1^), or vehicle control (0.1% DMSO) for 72 h. Growth rates were calculated using an exponential growth equation and were plotted against HHQ concentration to determine IC_50_ at 72 h post exposure as described previously ^15^.

To examine the long-term impacts of HHQ, triplicate flasks of 30 ml cultures of *E. huxleyi* (∼ 50,000 cells ml^-1^) were exposed to either 1 or 100 ng ml^-1^ HHQ, or a vehicle (0.1% DMSO) control. The experiment was sampled daily for 21 days to monitor *E. huxleyi* abundance, forward scatter, side scatter, red fluorescence, and photosynthetic efficiency (Fv/Fm). Fv/Fm was measured using a Fluorescence Induction and Relaxation (FIRe) system (Satlantic). Samples were dark adapted for 30 min, and photosystem II kinetics were measured from the average of 10 iterations of an 80 µs single turnover event and 1000 ms of weak modulated light.

To measure recovery, after 24, 48, 72, 96 h, or 21 d of HHQ exposure, triplicate 2 ml aliquots of HHQ-exposed culture was transferred into 198 ml of fresh media, effectively diluting HHQ to 1 ng ml^-1^. The same dilution was made with the vehicle control treatment, and the experiment was sampled daily for *E. huxleyi* growth rate, forward scatter, side scatter, and red fluorescence.

To investigate viral infection dynamics, triplicate 50 ml cultures were prepared for the following treatments: *E. huxleyi* (∼ 40,000 cells ml^-1^) + vehicle control (0.1% DMSO), *E. huxleyi* + EhV 207 (3.2 x 10^6^ EhV ml^-1^), *E. huxleyi* + HHQ (100 ng ml^-1^), *E. huxleyi* + HHQ + EhV 207. Samples were taken daily to monitor *E. huxleyi* abundance.

For all growth experiments, excluding the IC_50_ calculation, significant differences between treatments were determined by comparing abundances over time using ANOVAR, followed by a Dunnett’s multiple comparisons test ^75^. All data was tested to ensure that it passed the assumptions for normality and sphericity prior to running the ANOVAR.

### Physiological Assays

Propidium iodide (PI) was used to quantitatively discriminate cell cycle stage in HHQ-exposed phytoplankton cultures over 122 h. Three replicate 2 L cultures were dosed with either 100 ng ml^-1^ HHQ or vehicle control (0.002% DMSO). Fixed cells were enumerated every 24 h via flow cytometry. Every 2 h, approximately 10^6^ cells were subsampled, pelleted, and washed twice via centrifugation at 3,214 x *g* for 15 minutes at 18 °C. The dry cell pellets were resuspended in 1 ml of ice-cold LCMS-grade methanol, transferred to microcentrifuge tubes and stored at -80 °C. To read, methanol-fixed cells were centrifuged at 16,000 x *g* for 10 minutes at 4 °C, methanol removed, and pellets were resuspended in 1 ml of 1X DPBS before re-pelleting by centrifugation at 16,000 x *g* for 10 minutes at 4 °C. The pellet was resuspended in 0.5 ml of FxCycle PI/RNAse solution (Thermo Fisher) and incubated for 30 minutes in the dark and then measured via flow cytometry (583/26 nm emission).

Diagnostic fluorescent dye assays were used to measure indicators of cell stress and programed cell death (PCD) following HHQ treatment. Intercellular reactive oxygen species (ROS), nitric oxide (NO) production, mitotoxicity, cytotoxicity, and quantification of caspase proteases and activity were measured in *E. huxleyi* (CCMP2090) following HHQ treatment (70 ng ml^-1^ or 100 ng ml^-1^) at various time points up to 72 h post-exposure. See SI Appendix, Supplementary Information Text for detailed protocols.

*E. huxleyi* DNA integrity was examined using a modified protocol for the Click-iT TUNEL Alexa Fluor 488 Imaging Assay kit (Thermo Fisher). Cells were assayed using the manufacturer protocol and were sampled after 46 h HHQ exposure, with tagged cells enumerated via flow cytometry (512/18 nm emission). See SI Appendix Supplementary Information text for detailed protocols.

### Transmission Electron Microscopy

Replicate 20 ml cultures of exponentially growing *E. huxleyi* (∼1 x 10^5^ cells ml^-1^) were exposed to either 100 ng ml^-1^ HHQ or vehicle control (0.2% DMSO) for 24 and 337 h (14 d). Samples were concentrated by filtration on a 0.45 µm polycarbonate filter and transitioned out of f/2 media via three sequential washes with 10 ml of 0.2 M sodium cacodylate buffer pH 7.4, then fixed in 2% glutaraldehyde in 0.2 M sodium cacodylate buffer, pH 7.4. Samples were post-fixed in 2.0% osmium tetroxide for 1 h at room temperature and rinsed in DH_2_O prior to *en bloc* staining with 2% uranyl acetate. After dehydration through a graded ethanol series, the cells were infiltrated and embedded in Embed-812 (Electron Microscopy Sciences). Thin sections were stained with uranyl acetate and lead citrate and examined with a JEOL 1010 electron microscope fitted with at Hamamatsu digital camera and AMT Advantage NanoSprint500 software.

### Transcriptomic and Proteomic Analysis

A large-scale culturing experiment was performed with axenic *E. huxleyi* (CCMP2090) treated with either three concentrations of HHQ (1 ng ml^-1^, 10 ng ml^-1^, 100 ng ml^-1^) or vehicle control (0.002% DMSO) for 72 h. Following HHQ/DMSO exposure, 400 ml subsamples were taken from each quadruplicate 2 L bottle at both 24 and 72 h for total RNA isolation and an additional 1200 ml subsample was taken at 72 h for total protein isolation. Total RNA and protein were isolated and quantified as described in SI Appendix, Supplementary Information Text.

For RNA-seq analysis, the KAPA Stranded mRNA-Seq library preparation kit (Kapa Biosystems) was used to prepare library samples and sequenced on the NextSeq platform (Illumina) to generate 75 bp paired-end reads. Low-quality reads and adaptor sequences were trimmed using Trimmomatic (V0.38; ^76^). Transcript abundances were determined using *Salmon* (V0.12.0; ^77^) and the Ensembl ^78^ gene predictions for *E. huxleyi* CCMP1516 (the non-axenic form of CCMP2090; ftp://ftp.ensemblgenomes.org/pub/protists/release-41/fasta/emiliania_huxleyi/cdna/) as a transcript target index (k-mer size = 23). Normalization and determination of significantly differentially abundant transcripts was preformed using the DESeq2 R package (V1.22.1; ^79^). Tests for differential expression were carried out with the Wald test using a negative binomial generalized linear model. Logarithmic fold change (LFC) estimates were shrunken using the apeglm package (V1.6.0; ^80^) within DESeq2. Resulting *p* values were adjusted using the Benjamini-Hochberg (BH) procedure (see SI Appendix, Supplementary Information Text).

For proteomic analysis, proteins were solubilized in urea, reduced, alkylated, and trypsin digested following ^81^. Resulting peptides samples were desalted with a mini-centrifugal C18 column following manufacturer’s instructions (Nest Group). Peptides were chromatographically separated (precolumn: 3 cm, 100 μm i.d.; analytical column: 30 cm x 75 μm i.d; resin: 3 μm C18-AQ) with a nanoAcquity UPLC System (2–35% ACN, 0.1% v/v formic acid; 250 nl min-1, 90 minute) directly inline with a Fusion Lumos Orbitrap Tribrid mass spectrometer (Thermo Fisher Scientific) operated in data independent acquisition mode (DIA) following methods in ^82^. To generate a peptide spectral library, 1 µg of a pooled sample containing equal parts from each peptide digest was analyzed with six gas phase fractions covering 400-1000 m/z in 100 m/z increments (4 m/z staggered MS2 windows, 2m/z overlap). Each bioreplicate was then quantified in single DIA analyses (MS1: 400-1000 m/z; 8 m/z staggered MS2 windows, 4m/z overlap).

In order to generate absolute abundance measurements of detected proteins, raw MS data files were processed using msconvert (ProteoWizard) for demultiplexing and peak picking. EncyclopeDIA (V0.7.4) was used to 1) search resulting fragmentation spectra against the UniProt E. huxleyi CCMP1516 protein and contaminant database (10.0 ppm precursor, fragment, and library tolerances), 2) provide peptide-level area under the curve (AUC) data, and 3) generate quantitative reports of identified peptides and proteins for each HHQ MS experiment (1% false discovery rate). Significant changes (p < 0.05) in protein abundances between HHQ treatment and vehicle control were calculated as log2 fold-change between treatments. Complete details of protein sample preparations, chromatographic separations, mass spectrometry detection and quantification can be found in SI Appendix, Supplementary Information Text.

### PARP Inhibition and Homology Modelling

To examine the impact of alkylquinolone exposure on mammalian PARP activity, an inhibition assay was performed using the PARP Universal Colorimetric Assay Kit (R&D systems) according to the manufacturer instructions. Human PARP enzyme (0.5 U) was exposed to 50 µM HHQ, 50 µM PQS, or vehicle control (0.25% DMSO) for 15 min prior to the addition of a PARP activity buffer. See SI Appendix Supplementary Information text for a detailed protocol.

The *E. huxleyi* sequence XP_005783504.1 was aligned to the Protein Data Bank (PDB) database to determine the closest structural homolog with a small molecular inhibitor veliparib in the active site that could lend insight into HHQ binding.

### Exploring the Biogeography of HHQ Biosynthesis Genes

A conserved gene locus encoding genes involved in alkylquinolone biosynthetic pathway were previously identified from *Pseudoalteromonas piscicida* (A757) ^15^ and used to search the Ocean Gene Atlas web-based platform to explore the biogeography of genes with homology to pqsABCDE operon responsible for HHQ synthesis. Protein sequences from *P. piscicida* (A757) (Genbank Accession numbers: KT879191–KT879199) were searched against the Ocean Microbial Reference Gene Catalog (OM-RGC, version 1) using BLASTp search tools with an initial customized e-value threshold of 1 x 10^−10^.

### Detection of HHQ in Environmental Samples

Seawater samples were collected along a cruise track from Manta, Ecuador to Tahiti from October to December 2013 (US GEOTRACES EPZT GP16) as described previously ^83^. Briefly, seawater was collected at 3m depth by a tow-fish and pumped at a flow rate of 250 ml min^-1^ through a 0.2 µm filter and a polytetrafluoroethylene column packed with 20 g of polystyrene resin (Bondesil ENV; Agilent). Each sample represents an integrated average of 400-600 L of water across a wide region. Samples were frozen onboard at -20 °C. Prior to analysis, thawed columns were rinsed with 500 ml of 18.2 MΩ cm ultra-high purity water (qH_2_O) and eluted with 250 ml of LCMS grade methanol. The extracts were concentrated by rotary evaporation and brought up in a final volume of 6 ml of qH_2_O that was stored at -20°C. The organic extracts were separated by high pressure liquid chromatography (Dionex Ultimate 3000) coupled to an Orbitrap Fusion MS (Thermo Scientific), with specific methodology found in the SI Appendix, Supplementary Information Text.

## Acknowledgements

We acknowledge the support from the Electron Microscopy Resource Laboratory at the University of Pennsylvania for TEM sample processing. We thank Vinayak Agarwal for homology modeling support and discussions, Kay Bidle for viral culture, Bradley Moore for tetrabromopyrrole, and Katie Barott for flow cytometry support. We thank members of the Whalen Laboratory including Ellysia Overton, Yongjie Gao, Carlotta Pazzi, Megan Coolahan, Shreya Kishore, Lucy Zhao for assistance in phytoplankton sampling for RNA and protein isolation and constructive discussions. We thank the Georgia Genomics and Bioinformatics Core facility for RNA sequencing. Funding for this work was supported by the NSF grant (OCE-1657808) awarded to K.E.W. and E.L.H. K.E.W. was also supported by a Faculty Research Grant from Haverford College as well as funding from the Koshland Integrated Natural Science Center and Green Fund at Haverford College. E.L.H. was also supported by a Sloan Foundation Research Fellowship. B.L.N was supported by NSF grant (OCE-1633939). M.C.M. was supported by a NIH training grant (T32 HG000035). Mass spectrometry was partially supported by the University of Washington Proteomics Resource (UWPR95794). D.R. was supported by funding through the Gordon and Betty Moore Foundation (Grant 6000), the Simons Collaboration for Ocean Processes and Ecology Grant (329108), and an NSF grant (OCE-1736280). R.B. was supported by an NSF Graduate Research Fellowship and an NSF grant (OCE-1829761).

## Data Deposition Statement

Sequences from this study have been deposited in the Gene Expression Omnibus and are accessible through GEO Series accession number GSE131846 (https://www.ncbi.nlm.nih.gov/geo/query/acc.cgi?acc=GSE131846). The raw mass spectrometry proteomics data and subsequent spectral libraries have been deposited to the ProteomeXchange Consortium via the PRIDE partner repository (https://www.ebi.ac.uk/pride/archive/projects/PXD011560).

