## SUPPLEMENTAL INFORMATION for "Bacterial quorum sensing signal arrests phytoplankton cell division and protects against virus-induced mortality"

This PDF file includes:

- Supplementary text
- SI References
- Figures S1 to S18
- Tables S1 and S2

Other supplementary materials for this manuscript include the following:

- Datasets S1 and S2

### Supplementary Information Text

#### Diagnostic biochemical assays assessing PCD/ROS response.

Cellular stress following HHQ treatment was examined via diagnostic fluorescent dye assays. Intracellular reactive oxygen species (ROS) production was measured by exposing 200 µl of exponentially growing *E. huxleyi* (2090) culture to 5 µM of CM-$H_2$DCFDA stain at 18 °C for 90 min in the dark. In triplicate, vehicle control or HHQ (100 ng $ml^{-1}$) was added to each sample well and ROS production was measured on the flow cytometer (Guava, Millipore) in 30 min intervals at 512/18 nm. *E. huxleyi* cultures exposed to 100 or 500 nM tetrabromopyrrole (TBP) for 1 and 2 hours served as a positive control for ROS production (1).

Intracellular nitric oxide production was measured by exposing 200 µl of exponentially growing *E. huxleyi* (2090) culture to 10 µM of DAF-FM diacetate at 18 °C for 60 min in the dark. In triplicate, vehicle control or HHQ (100 ng $ml^{-1}$) was added and NO production was immediately monitored on the flow cytometer at 512/18 nm. Additional cultures were monitored for the presence of NO at 24 and 72 h post exposure as described above.

Both mitotoxicity and cytotoxicity following HHQ exposure was assessed using the HCS Mitochondrial Health Kit (Thermofisher) according to manufacturer's instructions. The MitoHealth stain detects changes in mitochondrial membrane potential, whereupon the reagent accumulates in active mitochondria while the stain Image-iT® DEAD$^{TM}$ Green detects cytotoxicity. To 200 µl of an exponentially growing *E. huxleyi* (2090) culture, 80 µl of Cell Staining Solution containing MitoHealth Stain and Image-iT® DEAD$^{TM}$ Green was added to each sample and incubated in the dark at 18 °C for 30 min. Triplicate samples were dosed with vehicle control or 100 ng $ml^{-1}$ HHQ and monitored at 512/18 nm and 575/25 nm on the flow cytometer for 24 h intervals. In a subsequent assay, the Image-iT® DEAD$^{TM}$ Green assay was repeated on triplicate *E. huxleyi* (2090) cultures exposed to vehicle control (DMSO) or 100 ng $ml^{-1}$ HHQ for 72 h and either 100 nM or 500 nM of TBP for 4.5 h, used as a positive control, as described above.

The induction of active caspase proteases was assessed using the CaspACE™ FITC-VAD-FMK In Situ Marker (Promega). Single replicates of 20 ml *E. huxleyi* cultures (~25,000 cells $ml^{-1}$) were exposed to either vehicle control or 100 ng $ml^{-1}$ HHQ. After 23 and 71 h, 400 µl of each culture was collected and stained with 10 µM of CaspACE dye for 1 h at 18 ºC in the dark before monitoring at 512/18 nm via the flow cytometer.

For each of the experiments above, significant differences between treatments were determined by comparing abundances over time using a repeated measures analysis of variance (ANOVAR), followed by Dunnett's multiple comparisons test.

Caspase activity following HHQ exposure was measured using the Caspase-8/FLICE Fluorometric Assay kit (BioVision). Duplicate 2 L batch cultures were exposed to either 100 ng $ml^{-1}$ HHQ or vehicle control (0.014% DMSO) for 72 h. At 72 h, 1.2 L of each culture was collected via centrifugation in acid-washed polypropylene bottles at 13,764 x *g* for 10 minutes at 4 °C to form a cell pellet that was transferred to a sterile 1.5 ml centrifuge tube. The pellet was centrifuged at 14,000 x *g* for 10 min at 4 °C. Supernatants were removed and dry cell pellets were flash frozen in liquid nitrogen prior to storage at -80 °C. Cell pellets were thawed on ice, resuspended in 1X Lysis Buffer (BioVision), lysed via sonication, and clarified via centrifugation at 12,000 x *g* for 10 min at 4 °C. Protein content of the clarified supernatants were measured using the Pierce Coomassie Plus Standard Microplate protocol (Thermo-Fisher). Triplicate samples of 100 µg of each protein lysate were added to a black-sided, clear-bottomed microplate and volume was raised to 100 µl with additional lysis buffer. Additional wells were prepared with 100 µl of lysis buffer or lysis buffer containing varying concentrations of active Caspase-8 protein (0.33 U or 1 U), which served as positive and negative controls, respectively. The active Caspase-8 protein was also added to 100 µg of protein lysate from DMSO-exposed cultures at final concentrations of 0.33 U and 1 U to demonstrate that pigments present in the protein lysates did not completely mask the fluorescent signal of the assay. To quantify caspase activity, 50 µM of IETD-AFC fluorogenic dye was added to each well. Fluorescence (Ex 400 nm, Em 505 nm) was collected every 1 min for 2 h via a Spectramax M2e plate reader (Molecular Devices) and HHQ versus vehicle control wells were compared at each time point using Welch's approximate t-test.

**TUNEL assay.**

*E. huxleyi* DNA integrity was examined using a modified protocol for the Click-iT TUNEL Alexa Fluor 488 Imaging Assay kit (Thermo Fisher). Five replicate 20 ml cultures of exponentially growing *E. huxleyi* (~2 x $10^5$ cells $ml^{-1}$) were exposed to either 100 ng $ml^{-1}$ of HHQ or vehicle control (0.2%). After 46 h, quadruplicate samples were collected from each culture by condensing ~5 x $10^5$ cells on a 0.45 µm nylon filter, transitioned to 1X PBS, and fixed at 4% paraformaldehyde in 1X PBS for 45 mins in the dark. The fixative was removed by filtration and samples were resuspended in HPLC-grade methanol for 1 h. The methanol was removed and cells were washed with nuclease free water. Water was removed and all samples were pretreated with Terminal deoxynucleotidyl Transferase (TdT) reaction buffer for 10 minutes before incubation with the TUNEL reaction mixture containing TdT enzyme plus a dUTP label for 1 h at 37 ºC in the dark. The reaction mixture was removed and all samples were washed twice with 1X PBS supplemented with 3% Bovine Serum Albumin (BSA). Following the removal of the second wash, samples were incubated with Click-iT reaction cocktail for 30 min in the dark to attach the Alexa Fluor 488 tagged dUTP molecules. Samples were washed with 1X PBS with 3% BSA, resuspended in 1X PBS, and monitored for forward scatter, side scatter, and green (512/18 nm) fluorescence emission.

**Direct assessment of DNA strand breaks.**

To assess if HHQ directly is capable of inducing DNA strand breaks, Lambda DNA (510 ng) and *E. huxleyi* (CCMP2090) genomic DNA (510 ng) was exposed to 100 ng $ml^{-1}$ HHQ or DMSO vehicle control at 18°C for 24 h in the dark. Duplicate samples were applied to a 1% agarose gel and visualized by ethidium bromide.

**Large scale culturing experiment for transcriptomic and proteomic analysis.**

Four 2 L axenic *E. huxleyi* batch cultures in exponential phase were used to inoculate sixteen 2 L cultures at final cell concentrations of approximately 6.72 x $10^4$ cells $ml^{-1}$. These cultures were grown for 48 h to approximately 1 x $10^5$ cells $ml^{-1}$, before quadruplicate cultures were exposed to either 1 ng $ml^{-1}$, 10 ng $ml^{-1}$, or 100 ng $ml^{-1}$ concentrations of HHQ or vehicle control (final concentration 0.002% DMSO in all bottles). Fixed cells were enumerated every 24 h by flow cytometry. For all treatments, 400 ml subsamples were taken from each bottle at 72 h for protein isolation, and at both 24 and 72 h for total RNA isolation.

Biomass for isolation of total RNA was collected in acid-washed autoclaved polycarbonate bottles via centrifugation at 13,764 x *g* for 8 min at 4 $^{o}$C. Supernatant was removed, and cell pellets were resuspended in 2 - 10 ml sterile seawater before aliquoting into microcentrifuge tubes (1 ml per tube). Biomass was again collected via centrifugation at 9,600 x *g* for 5 min at 4 $^{o}$C, the supernatant was removed, and pellets were flash frozen in liquid nitrogen prior to storage at -80 $^{o}$C. Total sampling time never exceeded 65 min per sample. Biomass for protein isolation was collected in acid-washed autoclaved polycarbonate bottles via centrifugation at 13,764 x *g* for 8 min at 4 $^{o}$C. Supernatant was removed, and cell pellets were resuspended in 20 - 30 ml sterile seawater before a second centrifugation at 1,500 x *g* for 8 min at 4 $^{o}$C. The majority of supernatant was removed before a final centrifugation at 1,500 x *g* for 5 min at 4 $^{o}$C and removal of all supernatant prior to flash freezing in liquid nitrogen and storage at -80 $^{o}$C. Total sampling time never exceeded 75 min per sample.

**RNA processing and sequencing.**

Total RNA was extracted using the RNeasy Plus Mini Kit (Qiagen) following the manufacturer's recommendations using 350 µl RLT plus buffer and the optional centrifugation step (18,800 x *g* for 1 min) prior to elution with 30 µl RNase free water. Eluent was reapplied to the column membrane and incubated for 8 min at room temperature before a second elution. Total RNA was quantified using a NanoDrop 2000 spectrophotometer (Thermo Scientific).

Total RNA for transcriptomic analysis was normalized to 20 ng $\mu l^{-1}$ before assessing integrity using the Agilent 2100 Bioanalyzer System Plant RNA assay (Agilent Technologies). Samples were then prepared for sequencing at the Georgia Genomics and Bioinformatics Core facility using the KAPA Stranded mRNA-Seq library preparation kit with KAPA mRNA capture beads (Kapa Biosystems) and sequenced on the NextSeq platform (Illumina) to generate 75 bp paired-end reads. Data discussed in this publication have been deposited in NCBI's Gene Expression Omnibus - accessible through GEO Series accession number GSE131846 (http://www.ncbi.nlm.nih.gov/geo/query/GSE131846).

**Sequence analysis.**

Reads were conservatively trimmed to remove adaptors, low-complexity and low-quality sequence, and rRNA reads (including chloroplast and mitochondria rRNA) using Trimmomatic (V0.38) with a custom adapter file and the following settings: ILLUMINACLIP:2:30:10 LEADING:3 TRAILING:3 MAXINFO:40:0.5 MINLEN:50. Read quality was examined before and after trimming using FastQC (V0.11.8; https://www.bioinformatics.babraham.ac.uk/projects/fastqc/) and MultiQC (V1.6; (2)). Salmon tool (V0.12.0) was used to determine transcript abundances in quasi-mapping mode with default settings and the following flags: --validateMappings; –gcBias. Quantification results were examined using MultiQC and processed using tximport (V1.10.0; (3)). Transcript and gene IDs were linked using the general feature format file for *Emiliania huxleyi* CCMP1516 (ftp://ftp.ensemblgenomes.org/pub/protists/release-41/gff3/emiliania_huxleyi). After estimation of size factors to normalize for differences in library sequencing depth and gene dispersion estimation, tests for differential expression were carried out using DESeq2 (V1.22.1) for each pairwise comparison of interest. Data visualization, including principal component analyses, were performed directly within the DESeq2 package after the data set was made homoskedastic by a regularized-logarithm transformation.

**Protein extraction and detection.**

Frozen cell pellets were thawed on ice and resuspended in 6 M urea in 4 $^{o}$C 50 mM ammonium bicarbonate buffer. Cells were lysed using a sonicating probe for 10 rounds (4 $^{o}$C, 10 s, 0.7-0.9 W) with 30 s rests between rounds. Lysate was centrifuged (16,000 x g for 10 min, 4 °C) and protein content of supernatant was determined using the colorimetric bicinchoninic acid assay (BCA) assay (Thermo Fisher) following manufacturer's instructions for a 96-well plate on a Spectramax 190 microplate reader (Molecular Devices).

To digest proteins, 100 µg of protein lysate in 100 µl with 6 M urea in 50 mM ammonium bicarbonate buffer was reduced using Tris (2-carboxyethyl) phosphine hydrochloride (TCEP; 2.5 µl of 200 mM) after the addition of tris buffer (pH 8.8) (6.6 µl of 1.5 M stock) to maintain a pH of 8 prior to incubating the sample for 1 h at 37 °C. Samples were then reduced with iodoacetamide (IAA) (20 µl of 200 mM stock; 1 h, 20 $^{o}$C, in dark) and remaining IAA was quenched with dithiothreitol (DTT; 20 µl of 200 mM;1 h, 20 $^{o}$C, in dark). Samples were diluted with 800 µl of 25 mM ammonium bicarbonate buffer and 200 µl HPLC grade methanol. Proteins were digested with 5 µg trypsin (12 h, 20 $^{o}$C, in dark). Samples were acidified with 30-40 µl of 50 % formic acid to pH < 2 before evaporating to dryness in a centrivacuum. Peptide samples were stored at -80 °C until analysis.

Peptide samples were resuspended in 2% ACN and 0.1% formic acid and desalted using centrifugal BioPure SPN MIDI PROTO solid phase extraction C18 column (PROTO, 300Å; Nest Group) following manufacturer's instructions. Final peptide samples were evaporated to dryness using a speedvac and resuspended in 100 µl of 2% ACN and 0.1% formic acid to final concentrations of 1 µg µl$^{-1}$. Individual sample vials were prepared by mixing 10 µl of peptide sample with 1 µl of 500 fmol µl$^{-1}$ Peptide Retention Time Calibration (PRTC; ThermoFisher Scientific) and 19 µl of 2% ACN and 0.1% formic acid. A pooled peptide sample was prepared by combining 3 µl from each individual sample vial and mixed by vortexing. Quality control (QC) samples (ThermoFisher Scientific) were prepared by mixing 10 µl of PRTC (Pierce Peptide Retention Time Calibration Mixture; Thermofisher) with 30 µl of 0.2% ACN, resulting in a final solution of 50 fmol µl$^{-1}$ PRTC.

Five QC samples were analyzed to ensure chromatographic reproducibility and ion intensity stability before starting sample data collection. For each sample analysis, 1 µg of sample peptides with 50 fmol of PRTC were chromatographically separated by reverse-phase chromatography using a 30 cm long, 75 µm i.d., fused silica capillary column (New Objective PicoTip) packed with C18 beads (Reprosil-Pur C18-AQ 3 µm (Dr. Maisch GmBH, Ammerbuch, Germany) in line after a with a 3 cm long, 100 µm i.d. precolumn (C18-AQ 3 µm Dr. Maisch GmBH). Peptides were eluted using a linear acidified (formic acid, 0.1% v/v) water-acetonitrile gradient (2–35% acetonitrile; 90 minute). Mass spectrometry (MS) was performed on the Lumos, operated in data independent acquisition mode (DIA) The pooled peptide sample was analyzed using 6 narrow-window (100 m/z each) gas phase fractionations (GPFs) spanning 400-1000 m/z with 4 m/z overlapping MS2 windows with mass spectrometry settings set to 120,000 precursor resolution and 30,000 fragment resolution, automatic gain control (AGC) target of 4e5, 60 ms maximum ion inject time (IIT), Normalized Collision Energy (NCE) of 33, and +2H default charge state. Each MS2 window for the pooled sample experiments overlapped with the next window by 2 m/z, proving high resolution and effectively duplicate 2 m/z sampling throughout. The pooled sample was used to generate a complete peptide spectral library for *E. huxleyi*. For individual sample quantification, DIA data was collected on ions 400-1000 m/z using 8 m/z staggered MS2 windows with MS settings of 120,000 precursor resolution, 15,000 fragment resolution, AGC target of 4e5, max IIT of 20 ms. Each MS2 window for the individual sample DIA experiments overlapped with the adjacent window by 4 m/z. The order of the MS experiments was randomized, and the first sample was re-injected after all other experiments were completed to determine if a chromatography shift had occurred. Quality control (QC) peptide mixtures (Pierce mixed peptide PRTC standards) were analyzed every 12th injection to monitor chromatography and MS sensitivity. Skyline was used to determine that QC standard retention time and isotopic distribution did not deviate > 10% through all analyses.

**Proteomic data analysis.**

All MS data was interrogated against a peptide database generated from *E. huxleyi* CCMP1516 genome (downloaded from Uniprot; downloaded Oct. 5, 2018) appended with a list of 50 common contaminant peptides and the QC standards.[1] All raw MS data was first demultiplexed by msconvert (ProteoWizard). A chromatogram library was generated from the pooled sample analyses using XCorDIA software in EncyclopeDIA 0.7.4. This analysis on the pooled samples provided peptide detection, fragment refinement, and determination of peak boundaries, generating a library of all detected peptides with associated peak area under the curve (AUC) values (a measure of absolute abundance). The chromatogram library ELIB file was generated in XCorDIA by processing mzML pooled files as "Jobs" against the background *E. huxleyi* FASTA proteome (35,707 proteins). Parameters adjusted based on experiment-specific details with the Data Acquisition Type set as "Non-Overlapping DIA." EncyclopeDIA was then used to search the wide-window individual sample data with the same "Background" UniProt FASTA for *E. huxleyi*. mzML files of individual samples were loaded as "Jobs" and processed with the same parameters as set for XCorDIA to generate quantitative reports, BLIB files, and a comprehensive chromatogram library. Detailed settings included: Target/Decoy Approach: normal; Data Acquisition Type: Non-overlapping DIA; Enzyme: Trypsin; Fixed: C+57 (Carbamidomethyl); Variable: None; Fragmentation: CID/HCD (B/Y); Precursor Mass Tolerance: 10.0 ppm; Fragment Mass Tolerance: 10.0 ppm; Maximum Missed Cleavage: 2; Percolator Version: v3-01; Percolator FDR threshold: 0.01; # Quantitative Ions: 5; Minimum # Quantitative Ions: 1; # Cores: 8; Charge range: 1 to 4. Quantitative reports of the detected proteins and the absolute abundance of these proteins were generated.

AUC data for all detected proteins in the quantitative report was visualized via a heatmap (ggplot2 R package). Principal component (PC) analysis of AUC data was performed by adding a value of 1 to each peak area value in the quantitative report, completing a logarithmic transformation of the data, and generating a principal component plot of PC1 and PC2 (ggbiplot package in R).

To examine the differential expression in each treatment, normality was confirmed visually and with the Shapiro-Wilk test. Equality of variance between the AUC values of HHQ treatments could not be determined via the F-test. The F-test indicated that 445 proteins (8% of total proteins detected) in the 100 ng $ml^{-1}$, 328 proteins (6%) in the 10 ng $ml^{-1}$, and 432 (8%) in the 1 ng $ml^{-1}$ HHQ treatments had significantly unequal variances. Welch's approximate t-test was therefore used to compare the mean AUC of proteins in each HHQ treatment (1-, 10-, 100 ng $ml^{-1}$) to the mean AUC of the DMSO control proteins. Differentially expressed proteins were

[1] Common contaminants in proteomics include proteins present in dust such as human or animal keratin.

determined using two levels of significance (adj. $p < 0.1$ and adj. $p < 0.05$) to stay consistent with the analysis of the transcriptomic data.

The change in protein expression level between each treatment and the DMSO control were calculated as log2 fold-change using the equation below (i.e., value of 1 is a 2-fold difference, while a value of 3 is an 8-fold difference). Negative values represent decreased abundance of proteins in the treatment relative to DMSO control, while positive values represent increased abundance in the treatment relative to DMSO control.

$$\log_2\left(\frac{\text{Mean Peak Area of Treatment}}{\text{Mean Peak Area of Control}}\right) = \log_2 \text{FC}$$

As only the 100 ng $ml^{-1}$ HHQ treatment had >5 differentially expressed proteins at the chosen levels of significance, heatmaps displaying the log2 fold-change data of each treatment for the list of differentially expressed proteins in the 100 ng $ml^{-1}$ high treatment were generated.

Lists of all GO identification codes associated with various gene/protein functions were generated by searching the cellular component, biological process, and molecular function GO hierarchies for key words associated with selected pathways (Dataset 2) (4). For each pathway, a specificity threshold ID was manually chosen (Dataset 2) and all GO IDs below the threshold were compiled. These lists of GO IDs were then used to color the most stringent expression level scatterplot (adj. $p < 0.05$) by pathway; GO terms were compiled and used to color the scatterplot. The NCBI Protein BLAST tool was used to compare the amino acid sequence of selected *E. huxleyi* proteins to better annotated translated genomes of other organisms (5). The BLAST results (significance threshold: e-value < $1 \times 10^{-20}$) were used to determine the likely identity of these selected proteins through sequence homology and to validate the biological category associated with each list of proteins.

**PARP inhibition.**

To examine the impact of alkyl quinolone exposure on mammalian poly(ADP-ribose) polymerase (PARP) activity, an inhibition assay was performed using the PARP Universal Colorimetric Assay Kit (R&D systems). In a histone-coated strip-well plate, duplicate wells without inhibitor or without enzyme served as positive and negative controls, respectively. Four replicates of 0.5 U of mammalian PARP enzyme was pre-incubated with 50 µM HHQ, 50 µM PQS, or vehicle control (0.25% DMSO) for 15 min prior to the addition of a PARP activity buffer containing biotinylated-nicotinamide and activated DNA for 1 h in the dark. Wells were washed twice with 200 µL of 1X PBS + 0.1% Triton X-100, followed by two additional washes with 1X PBS before the addition of Strep-HRP for 1 h in the dark. The wells were washed as before and 50 µl of pre-warmed TACS-Sapphire colorimetric substrate was incubated for 15 minutes in the dark. The reaction was stopped and color stabilized by the addition of 50 µl of 0.2 M HCl. Absorbance at 450 nm was measured using Tecan Infinite M200 plate reader.

**Detection of HHQ in environmental samples.**

Organic seawater extracts were separated by high pressure liquid chromatography (Dionex Ultimate 3000) coupled to an Orbitrap Fusion MS (Thermo Scientific). Separations were performed on 50 µL injections on a C8 column (2.1 × 100 mm, 3 µm particle size; Hamilton) using a 50 min gradient from 10% to 90% methanol in water with 5 mM ammonium formate (LCMS grade, Sigma-Aldrich) as a buffer. The orbitrap was equipped with a heated electrospray ionization source with a capillary voltage of 3500 V; sheath, auxiliary, and sweep gas flow rates of 12, 6, and 2 (arbitrary units); and ion transfer tube and vaporizer temperatures of 300 and 75 °C. Mass spectra were collected in positive mode at 450K resolution. Fragmentation spectra were collected using a quadrupole isolation window of 1 m/z and high-energy collision-induced dissociation (HCD) energy of 35%. The limit of detection for HHQ using the described method was 0.18 ng $L^{-1}$.

**Figure S1.** Effect of HHQ and PQS on *E. huxleyi* growth. Dose-response curve of *E. huxleyi* (strain 2090, axenic) in response to (a) HHQ and (b) PQS. Each symbol represents the mean of three independent replicates ± standard deviation. An inhibitory concentration ($IC_{50}$) of 84.3 ng $ml^{-1}$ was calculated from the HHQ curve.

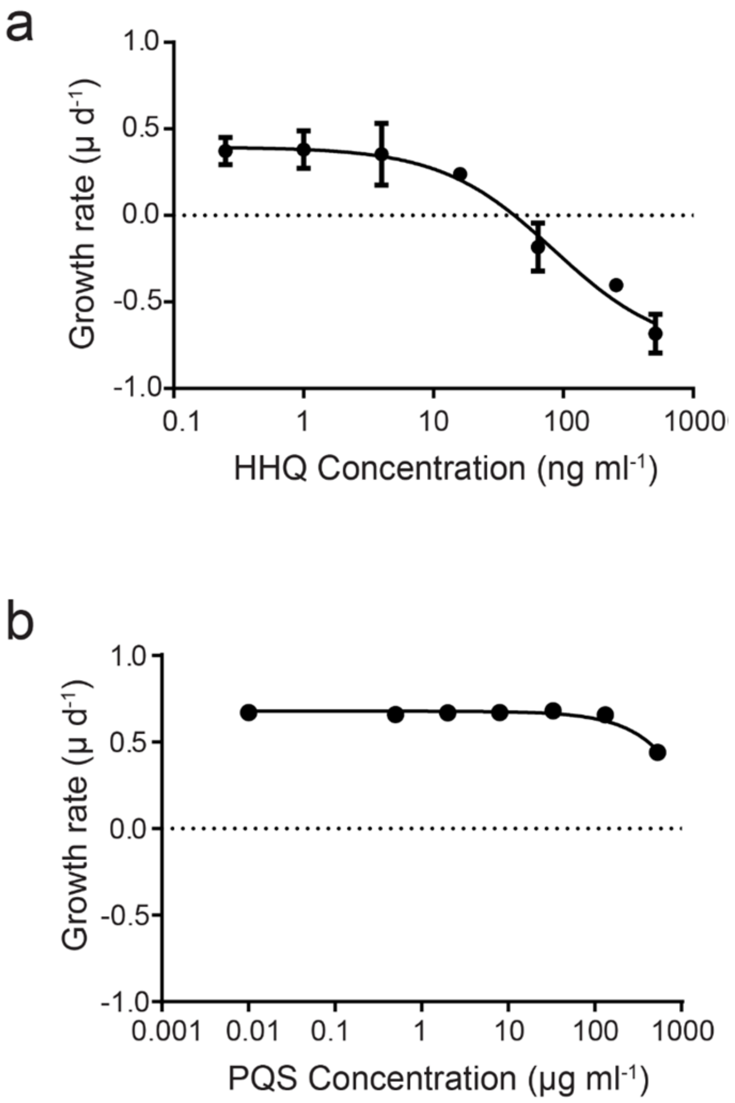

**Figure S2.** The mean side scatter (a) and photosynthetic efficiency (b) of *E. huxleyi* exposed to 1- (blue) and 100- (red) ng ml$^{-1}$ HHQ or a vehicle control (gray) over 504 h. Using a repeated measures ANOVA, it was determined that the 100 ng ml$^{-1}$ HHQ-exposed cells had a significantly higher side scatter relative to the vehicle control ($p < 0.05$), however no significant differences in photosynthetic efficiency were observed among the treatments. Each symbol represents the mean of three independent replicates ± standard deviation.

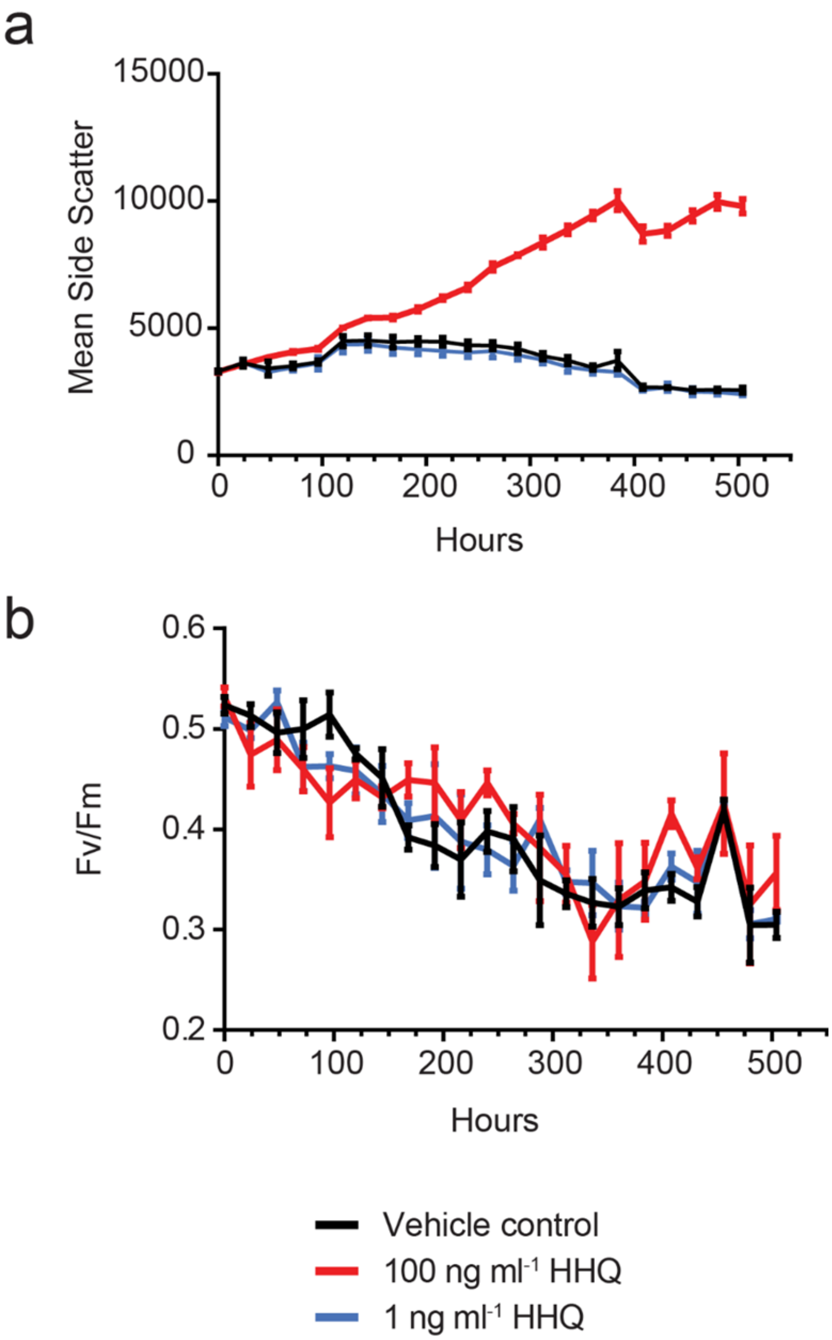

**Figure S3.** Composite TEM images of *E. huxleyi* cells (n = 9) exposed to vehicle control (DMSO) for 24 h. All nine micrographs show representative images of *E. huxleyi* cells exposed to DMSO for 24 h. In the first image, subcellular structures have been labeled for identification. Chloroplast (c), mitochondria (m), nucleus (n), pyrenoid (p), golgi (white arrow), vacuole (v). Scale bar = 500 nm.

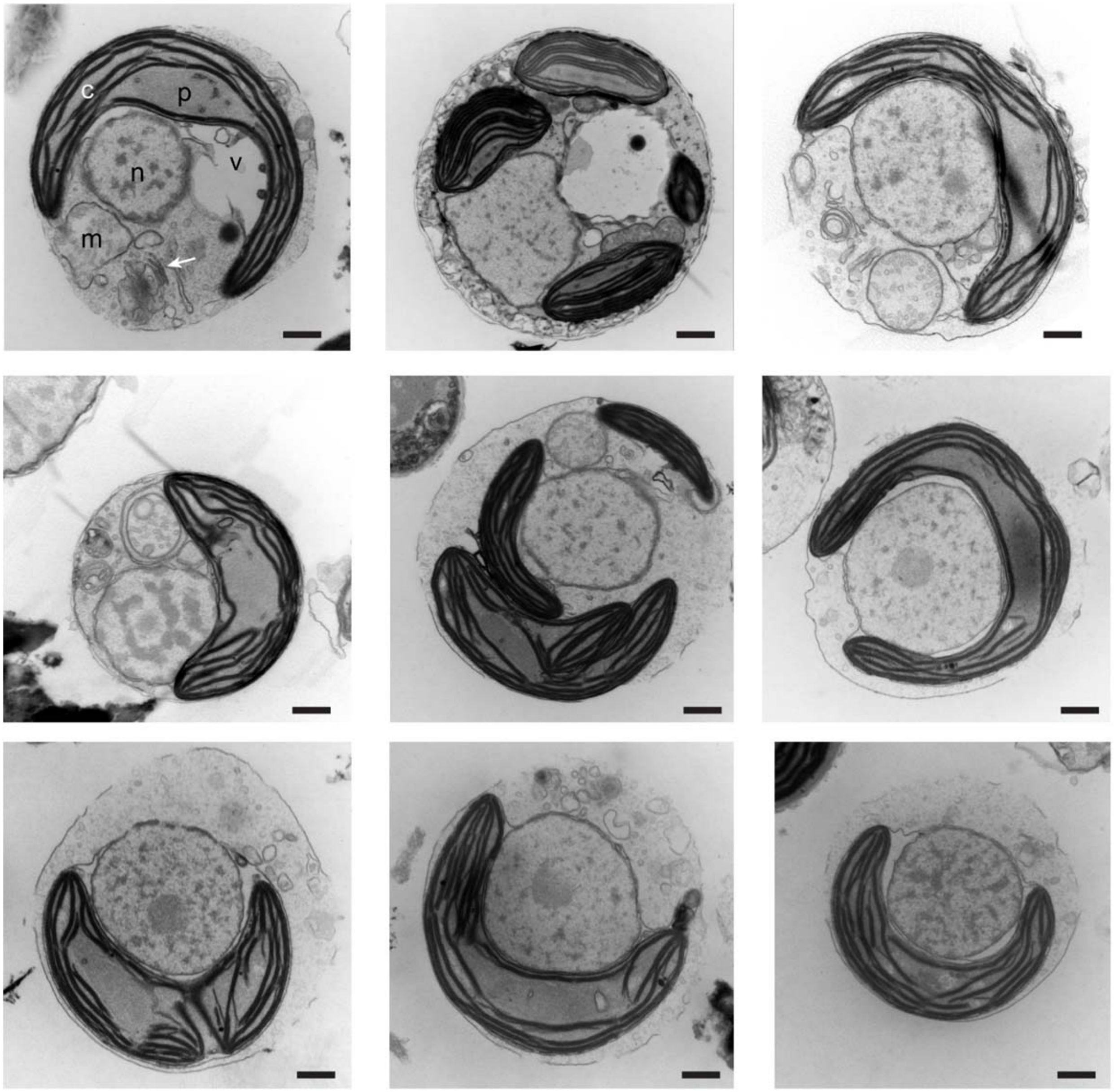

**Figure S4**. Composite TEM images of *E. huxleyi* cells (n = 9) exposed to 100 ng $ml^{-1}$ HHQ for 24 h. All nine micrographs show representative images of *E. huxleyi* cells exposed to HHQ for 24 h, highlighting a diversity of ultrastructure changes in comparison to vehicle control exposed cells (see Figure S3). Select micrographs have subcellular structures labeled for identification. Chloroplasts with distended thylakoid membranes (c), lipid droplet (l), mitochondria (m), nucleus (n), vesicles (arrow heads). Scale bar = 500 nm.

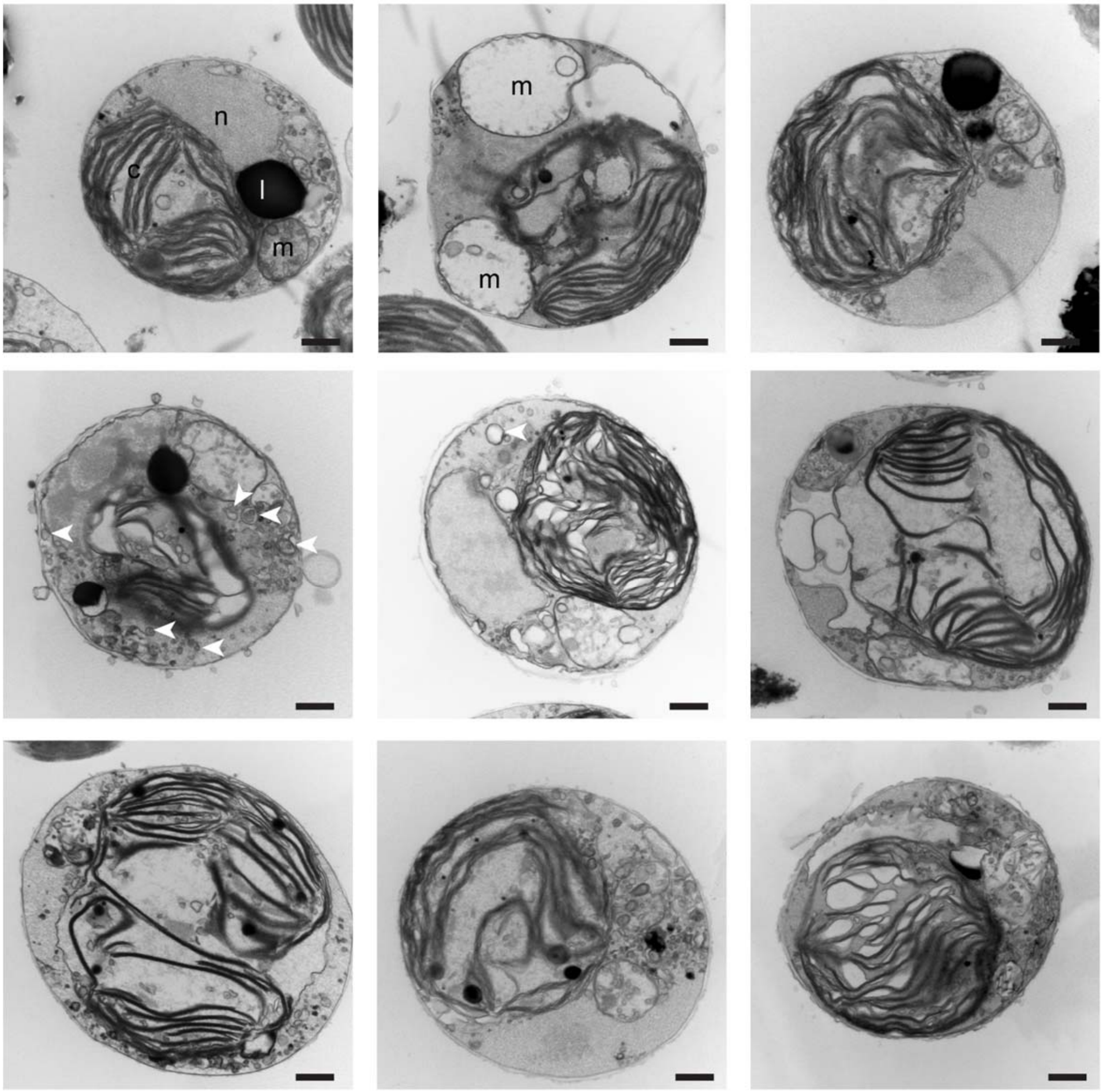

**Figure S5.** Composite TEM images of *E. huxleyi* cells (n = 12) exposed to 100 ng ml$^{-1}$ HHQ for 14 d. All twelve micrographs show representative images of *E. huxleyi* cells exposed to HHQ for 14 d, highlighting a diversity of ultrastructure changes in comparison to vehicle control exposed cells (see Figure S3). Select micrographs have subcellular structures labeled for identification. Chloroplasts with distended thylakoid membranes (c), golgi (g), lipid droplet (l), mitochondria (m), nucleus (n), nucleolus (nu), pyrenoid (p), vacuoles (v), vesicles (white arrow heads).

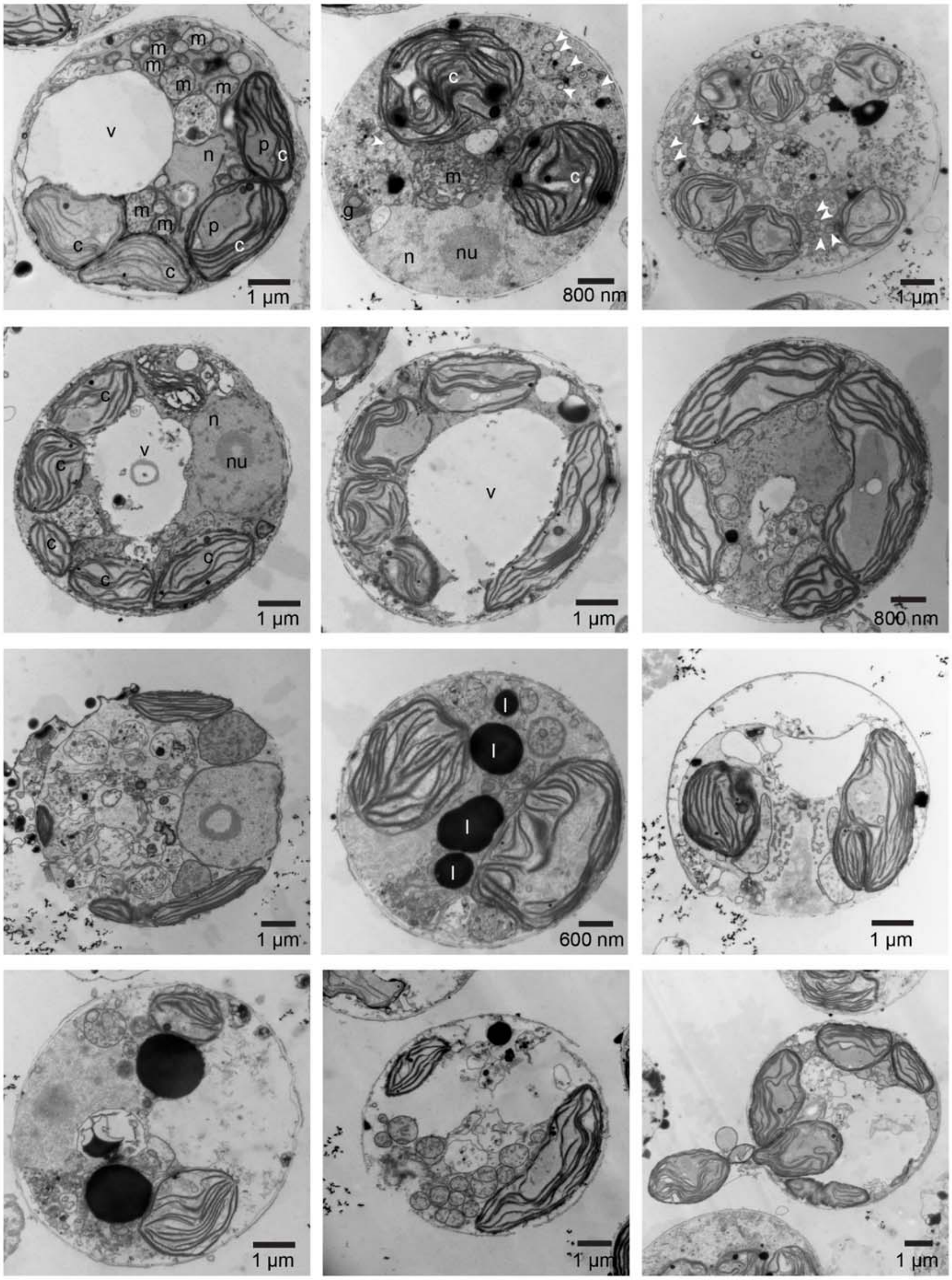

**Figure S6.** *E. huxleyi* cells exposed to 100 ng $ml^{-1}$ of HHQ exhibit an increase in relative chlorophyll fluorescence per cell relative to HHQ-insensitive strains of eukaryotic phytoplankton (i.e. *Dunaliella* and *Phaeodactylum*). Each symbol represents the mean of three independent replicates ± standard deviation. Data were fit to a second order polynomial equation using non-linear regression. Statistical significance using a t-test was determined using the Sidak-Bonferroni method with an alpha = 0.05.

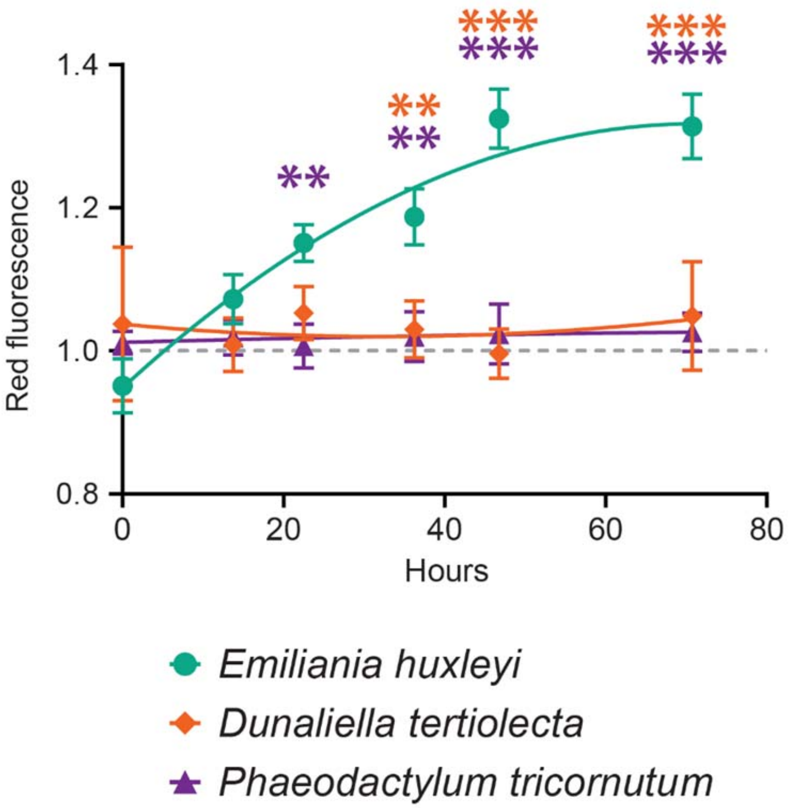

**Figure S7.** *E. huxleyi* cultures were examined for their ability to recover following exposure to 100 ng ml$^{-1}$ of HHQ at five time points extending up to 504 h. Following HHQ exposure, cultures were monitored for their ability to recover by measuring growth rate (a thru e; each point represents the exponential growth rate over the previous 24 h), red fluorescence (f thru j), and forward scatter (k thru o). Each symbol represents the mean of three independent replicates ± standard deviation. Significant differences between HHQ-exposed cells and the vehicle control were assessed using an ANOVAR ($p < 0.05$).

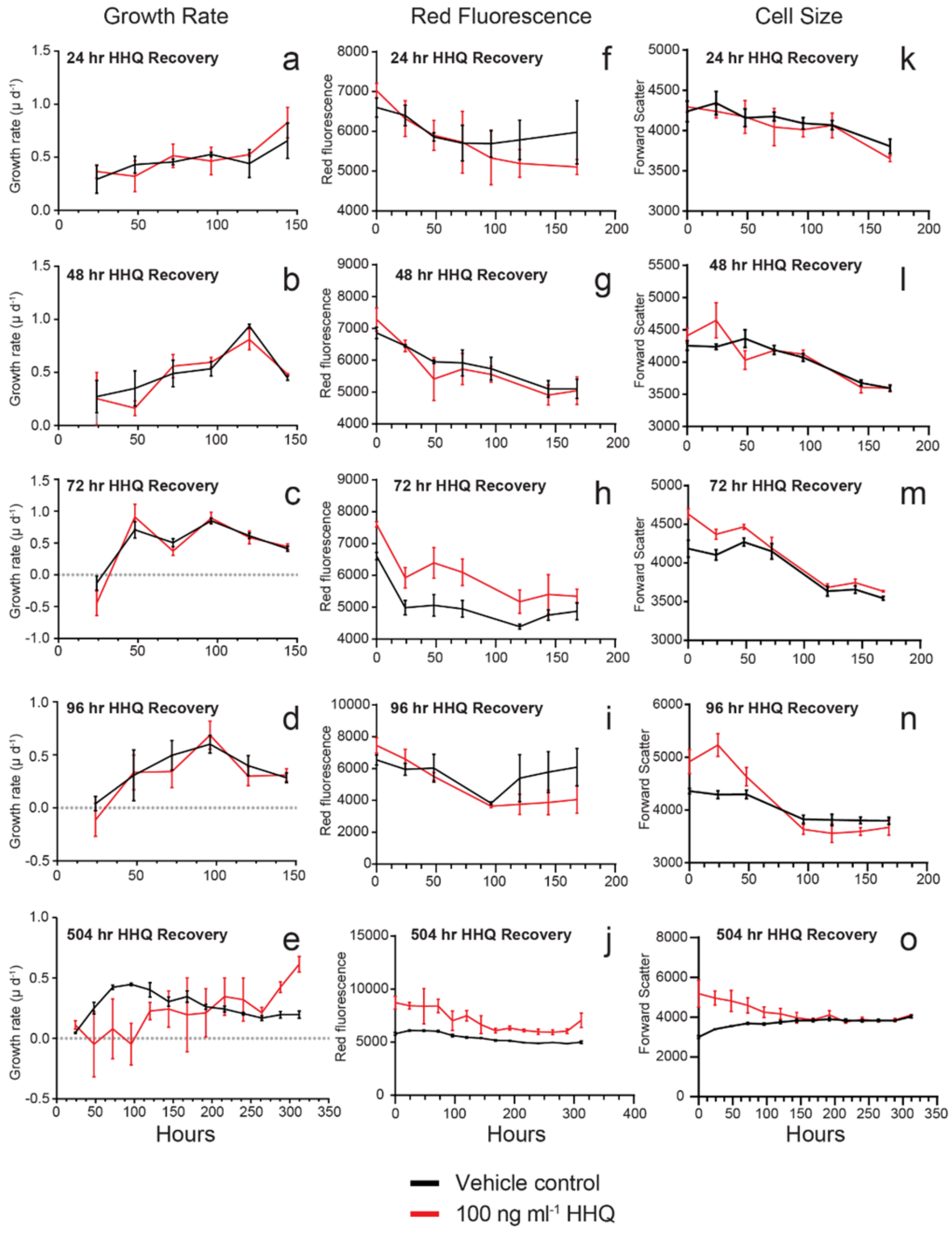

**Figure S8. (**a). Principal component analysis performed on the regularized-logarithm transformed transcriptomic read counts matrix. Each point represents an experimental sample with point color indicating treatment and shape indicating the time point. The first component explains 39% of variance and the second component explains 20%. (b). Principal component analysis performed on logarithm transformed proteomic data matrix (protein summed peptide peak area values +1). Each point represents an experimental sample with point color indicating treatment. The first component explains 29.8% of variance and the second component explains 16.2%. Note, protein was only collected at the 72 hr.

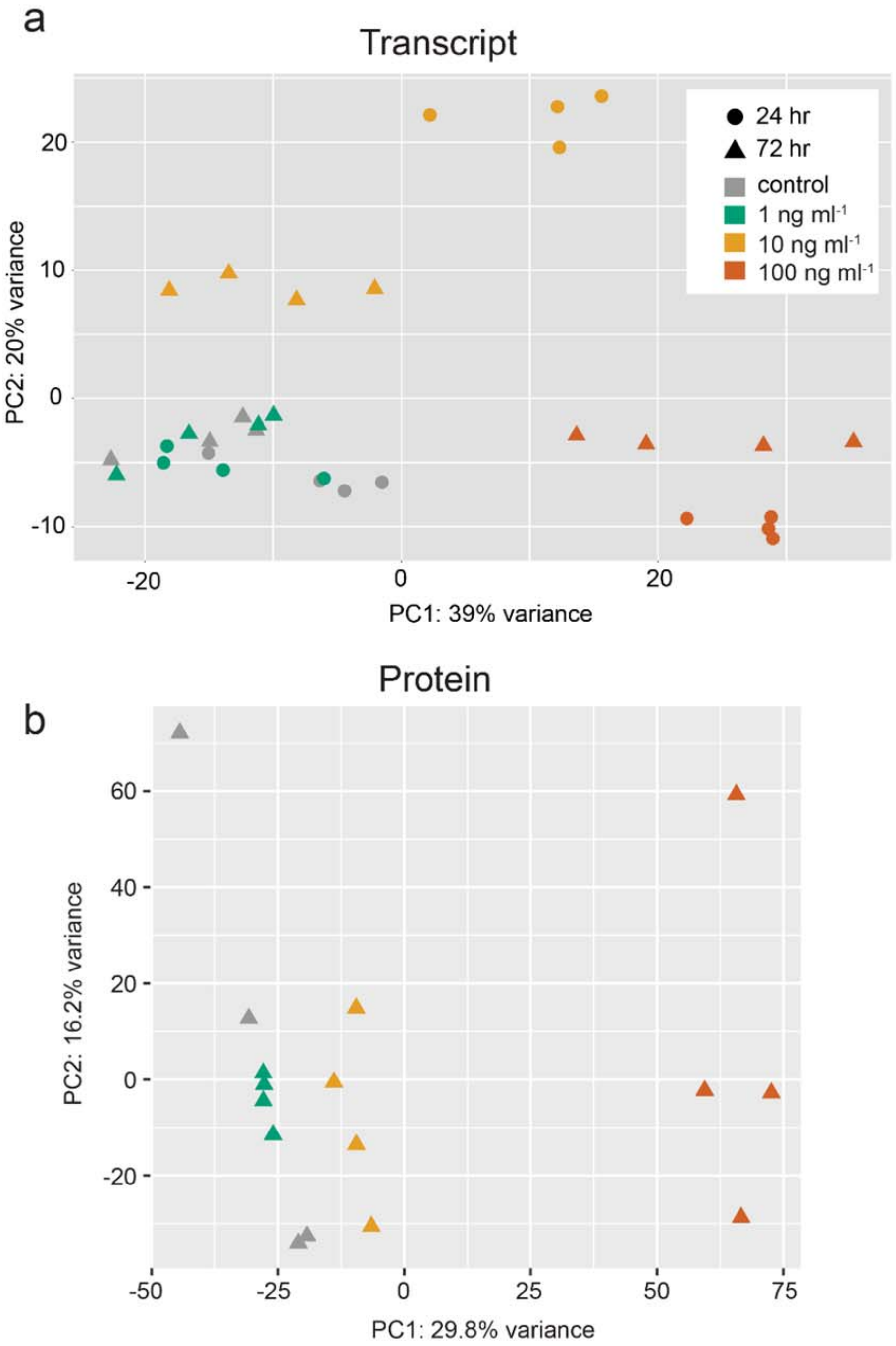

**Figure S9.** Hierarchical clustering analysis of the top 500 *E. huxleyi* transcripts exhibiting the greatest variance following 24 h (a) and 72 h (b) exposure to HHQ. HHQ treatment type (1-, 10-, 100 ng $ml^{-1}$, DMSO vehicle control) are shown in columns and genes are shown in rows. Coloring of the heatmap is dictated by the regularized-logarithm transformed counts for the top 500 transcripts that exhibited the greatest variance from their mean across the entire data set.

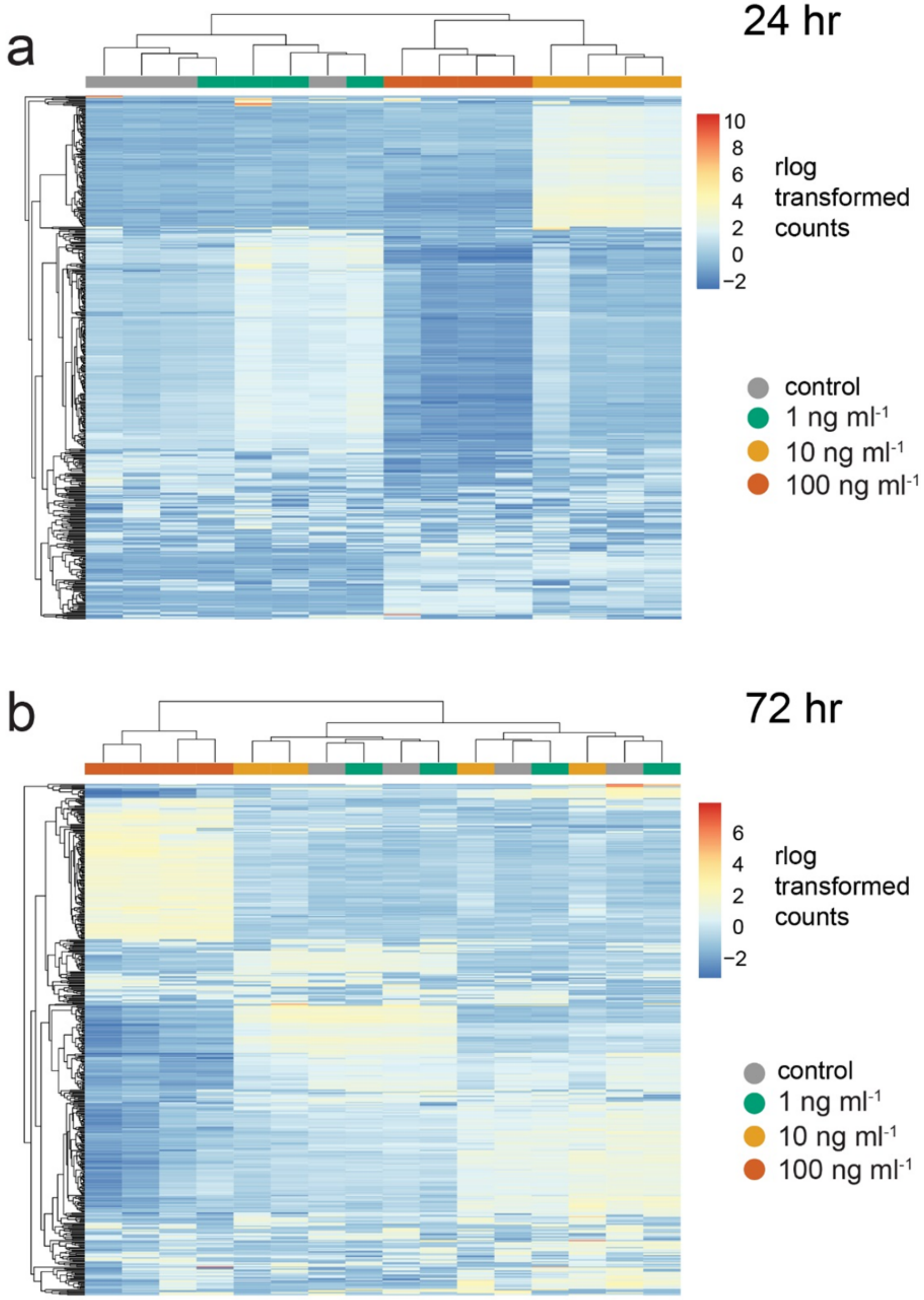

**Figure S10.** Hierarchical clustering of the 879 differentially expressed *E. huxleyi* proteins in each treatment compared to the vehicle control (Welch's approximate t-test, *q*-value < 0.05). Coloring is dictated by the $\log_2$ fold change in expression level of the HHQ treatment type (1-, 10-, 100 ng $ml^{-1}$) vs the vehicle control (DMSO). Treatment types are shown in columns with proteins shown in rows.

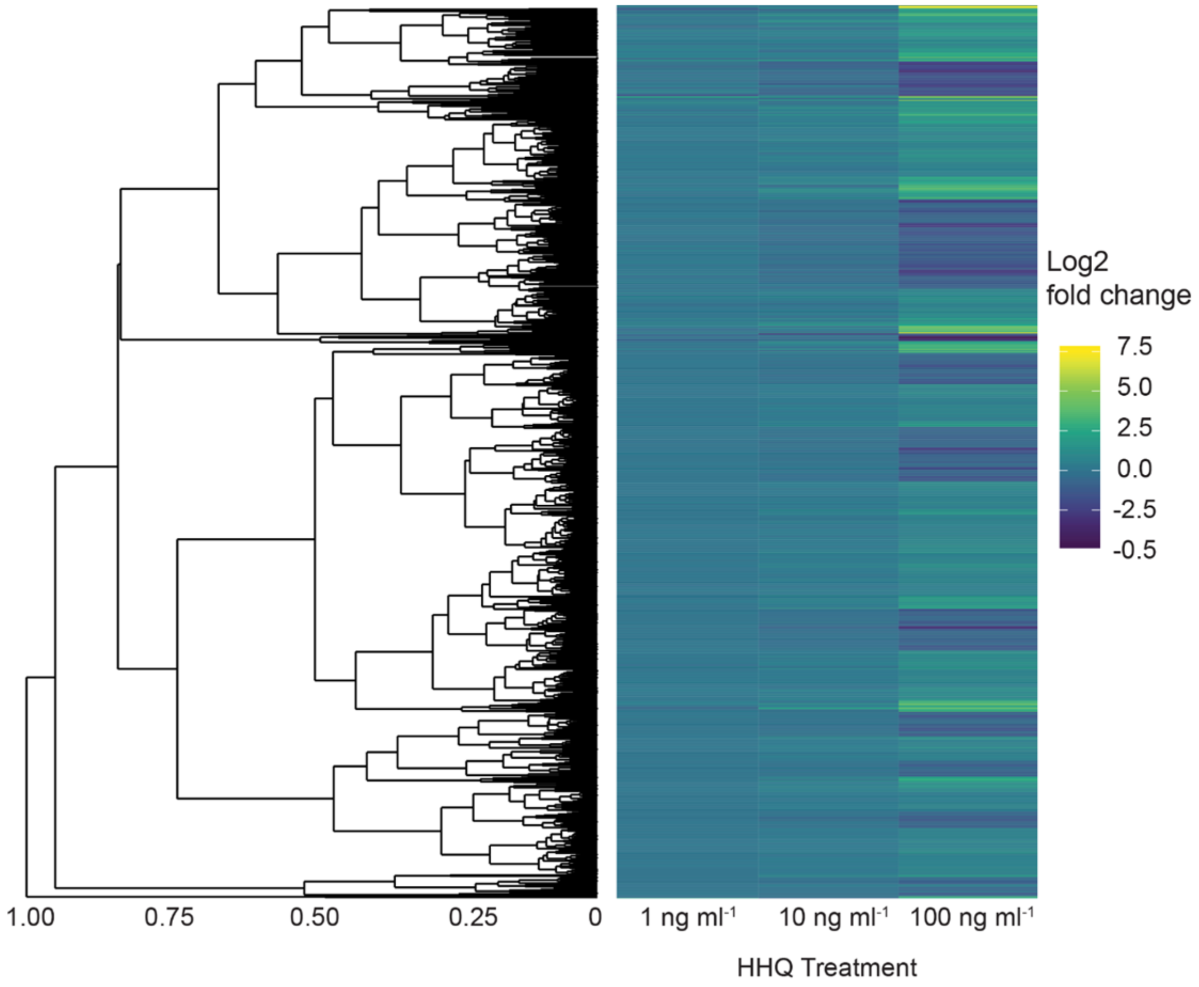

**Figure S11.** MA plots of transcriptional changes induced by HHQ treatments at 24 (a thru c) and 72 h (d thru f) post exposure. The log2 fold change for each HHQ treatment (1, 10, 100 ng $ml^{-1}$) compared to the vehicle control is plotted as a function of the average of the counts normalized by size factor. Genes are represented by dots and those with an adjusted p value below 0.05 are colored red. Log2 fold changes were shrunk prior to plotting using the apeglm method for shrinking coefficients (Zhu et al. 2018; Love et al. 2014).

Caption references
Zhu, A., Ibrahim, J.G., Love, M.I. (2018) Heavy-tailed prior distributions for sequence count data: removing the noise and preserving large differences. Bioinformatics. https://doi.org/10.1093/bioinformatics/bty895
Love MI, Huber W, Anders S (2014). “Moderated estimation of fold change and dispersion for RNA-seq data with DESeq2.” Genome Biology, 15, 550. doi: 10.1186/s13059-014-0550-8.

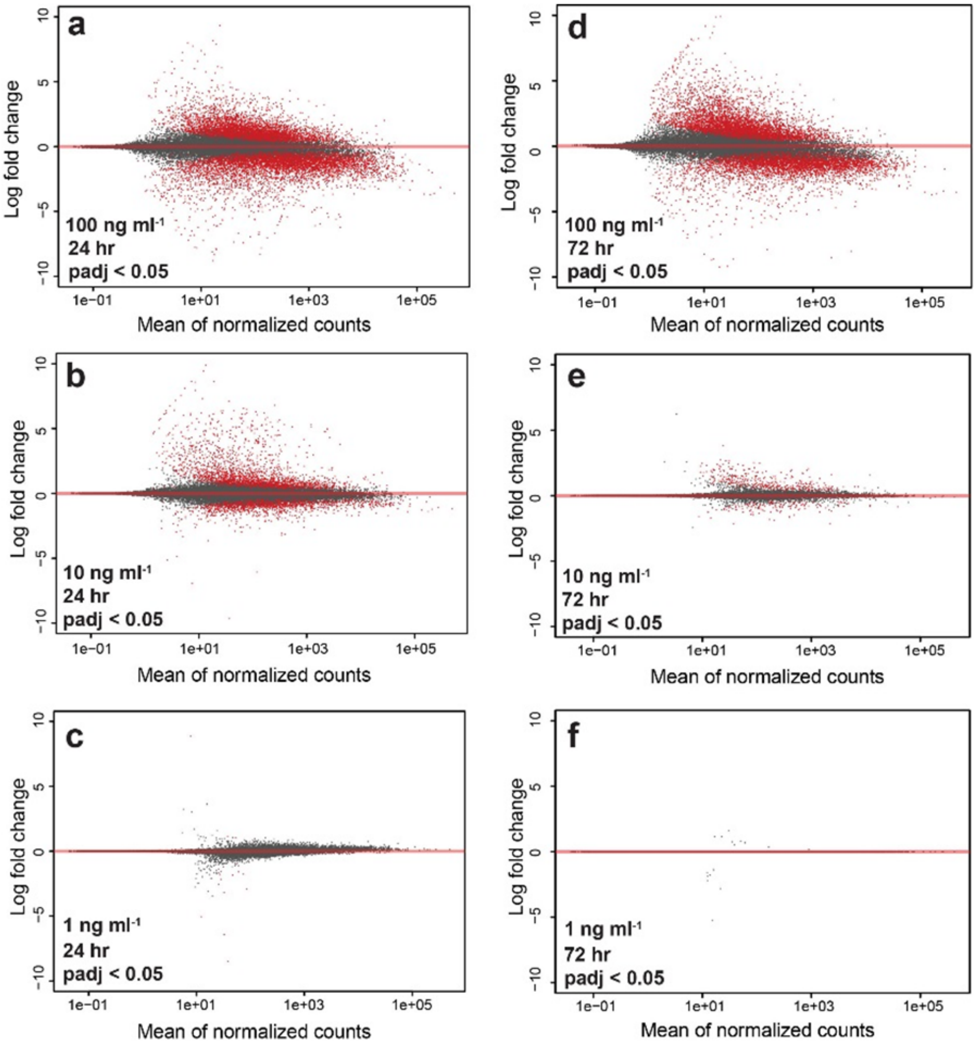

**Figure S12.** Volcano plots of proteomic changes induced via exposure to HHQ for 72 h. The $\log_2$ fold change of the protein abundance in the HHQ treatment (1, 10, 100 ng $ml^{-1}$; a, b, c, respectively) compared to the vehicle control is compared to the negative $\log_{10}$ transformed *q*-value (Benjamini-Hochberg method). Proteins are represented by dots and the dashed line indicates a value of 1.301, which represents a *q*-value of 0.05. All proteins that are significantly differentially abundant (*q*-value < 0.05) are colored red.

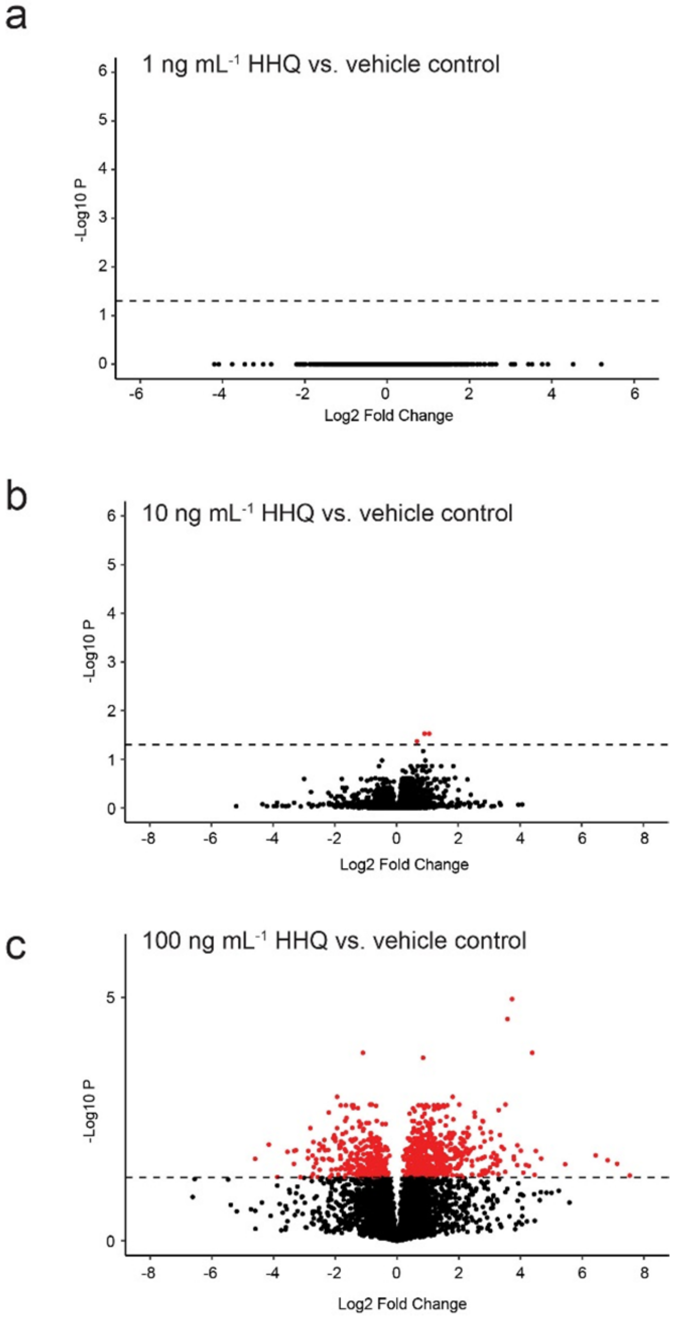

**Figure S13.** Following exposure to 100 ng ml$^{-1}$ of HHQ cells arrest in S-phase. (a) Density plots of the distribution of cells with varying DNA contents ranging from 2N (G1, yellow) to 4N (G2, blue). Cells with intermediate DNA content were denoted as S-phase (orange), as the genome replicated. Each plot represents the distribution of a single replicate for each treatment taken every 12 h. A dashed line was put at 1000 relative fluorescence unit (RFU) for reference. (b) Average cell abundance of triplicate cultures exposed to HHQ or the vehicle control over the course of the cell cycle analysis. Error bars represent the standard deviation of the average cell abundance for each treatment. The arrow indicates the point when HHQ or the vehicle control was added (T= 0 h).

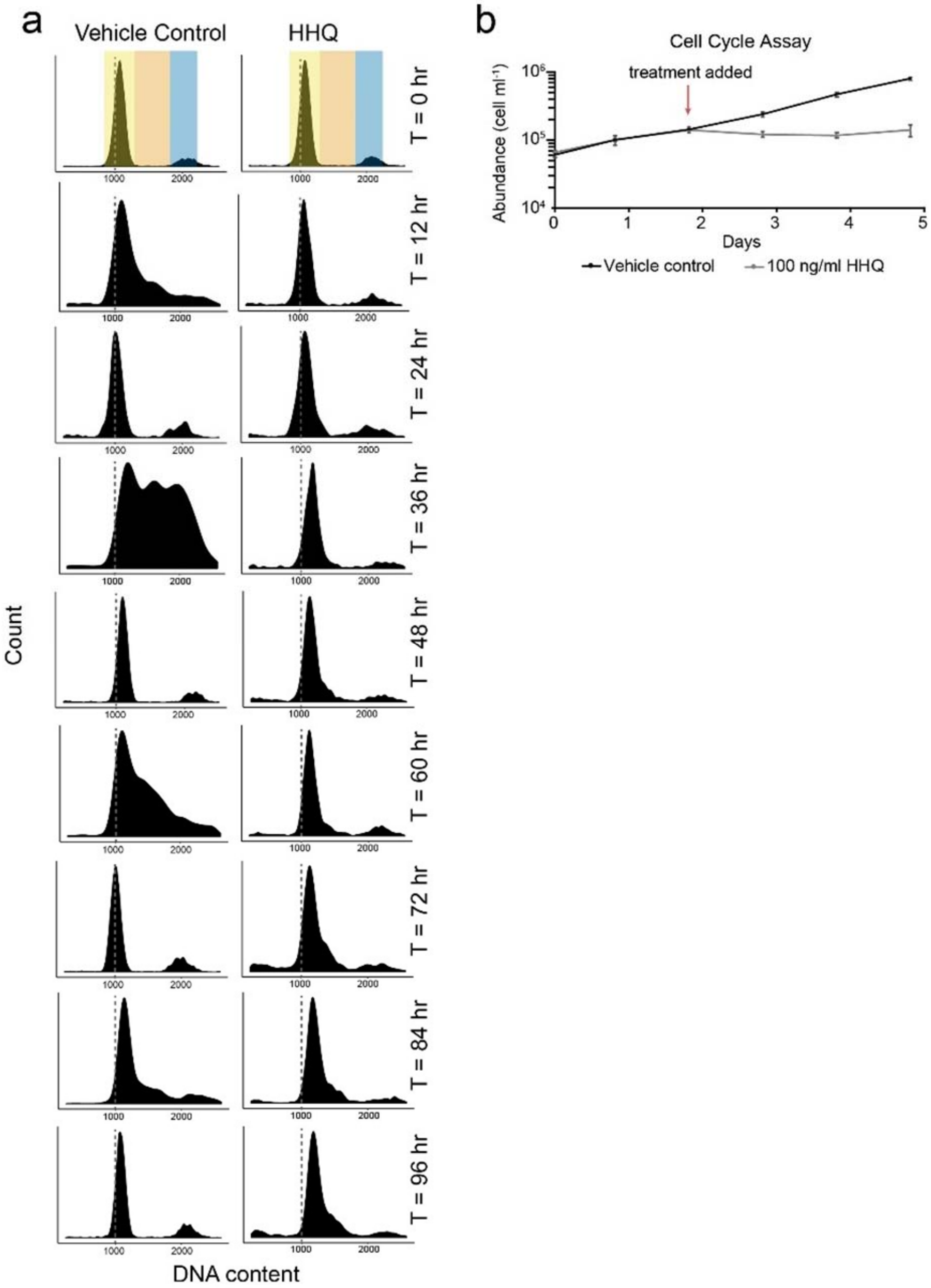

**Figure S14.** Diagnostic biochemical assays for (a) caspase activity from whole-cell lysates or purified caspase enzyme (comparison of protein – 72 hr DMSO versus protein – 72 hr HHQ; Welch's approximate t-test, *p*-value < 0.05; no significance observed), (b) the presence of active caspase proteases *in vivo* (CaspACE), (c) reactive nitrogen species (DAF-FM Diacetate), (d) reactive oxygen species (CM-$H_2$DCFDA) from DMSO or HHQ-exposed cultures from 1 to 72 h post-treatment, (e) reactive oxygen species (CM-$H_2$DCFDA) from 72 h DMSO or HHQ-exposed cultures that were spiked with the algicide tetrabromopyrrole (TBP, positive control) for 1 or 2 h, (f) apoptosis (Image-iT DEAD stain) from DMSO or HHQ-exposed cultures from 1 to 72 h post-treatment, (g) apoptosis (Image-iT DEAD stain) from 72 h DMSO or HHQ-exposed cultures that were spiked with TBP (positive control) for 4.5 h, and (f) mitochondrial membrane integrity (MitoHealth Stain). Bars represent the mean ± standard deviation of triplicate readings. Significance (part b, c, d, f, h) based on ANOVAR, Dunnett's multiple comparisons test, *p*-value < 0.05. Significance (part e, g) based on Student's t-test, *p*-value < 0.05. Cells exposed to HHQ were only significantly different from the control using the MitoHealth stain after 24 h of exposure. However, the two treatments were not significantly different from one another at all subsequent time points.

**Figure S15.** Exposure to 100 ng ml$^{-1}$ HHQ leads to DNA damage and disrupts the DDR response. (a) Cultures of *E. huxleyi* were exposed to 100 ng ml$^{-1}$ HHQ or the vehicle control (DMSO) for 46 h before pigments were removed and cells were stained using an *in vivo* TUNEL assay to detect presence of DNA ends, a proxy for DNA breaks. (b) Lambda DNA and genomic DNA isolated from *E. huxleyi* CCMP2090 was incubated with 100 ng ml$^{-1}$ HHQ or the volumetric equivalent of DMSO for 24 h at 18°C, and the presence of DNA strand breaks was assessed by agarose gel stained with ethidium bromide. (c) Inhibition of human PARP-1 enzyme by 50 µM HHQ and PQS. Percent PARP inhibition of was measured using a colorimetric assay by subtracting the ratio of absorbance values of quadruplicate wells containing HHQ or PQS,

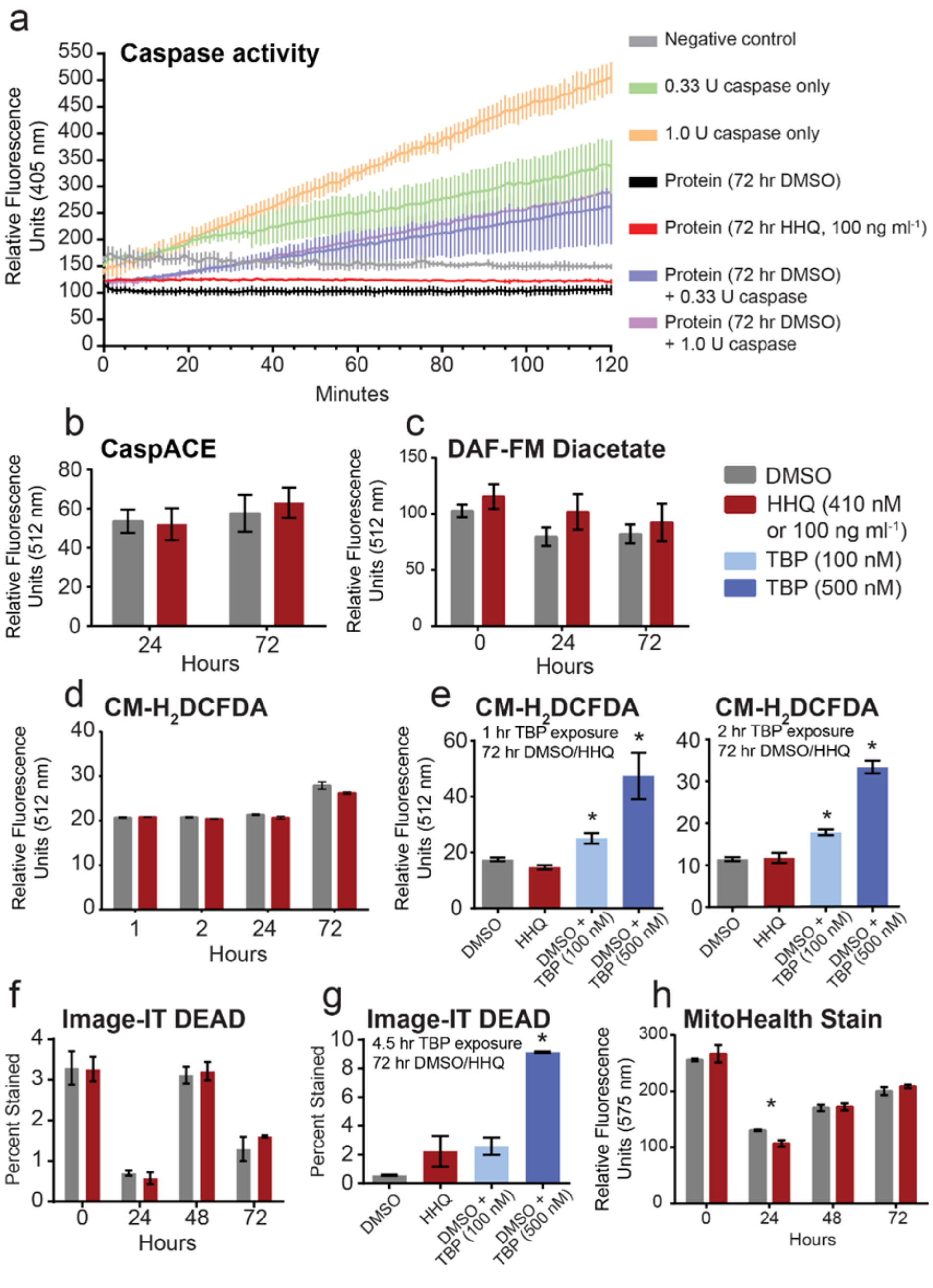

compared to the vehicle control, from 100 percent. Points represent individual replicates. Asterisks indicate a significant difference between the treatment and the vehicle control (Welch's approximate t-test, *p*-value < 0.05)

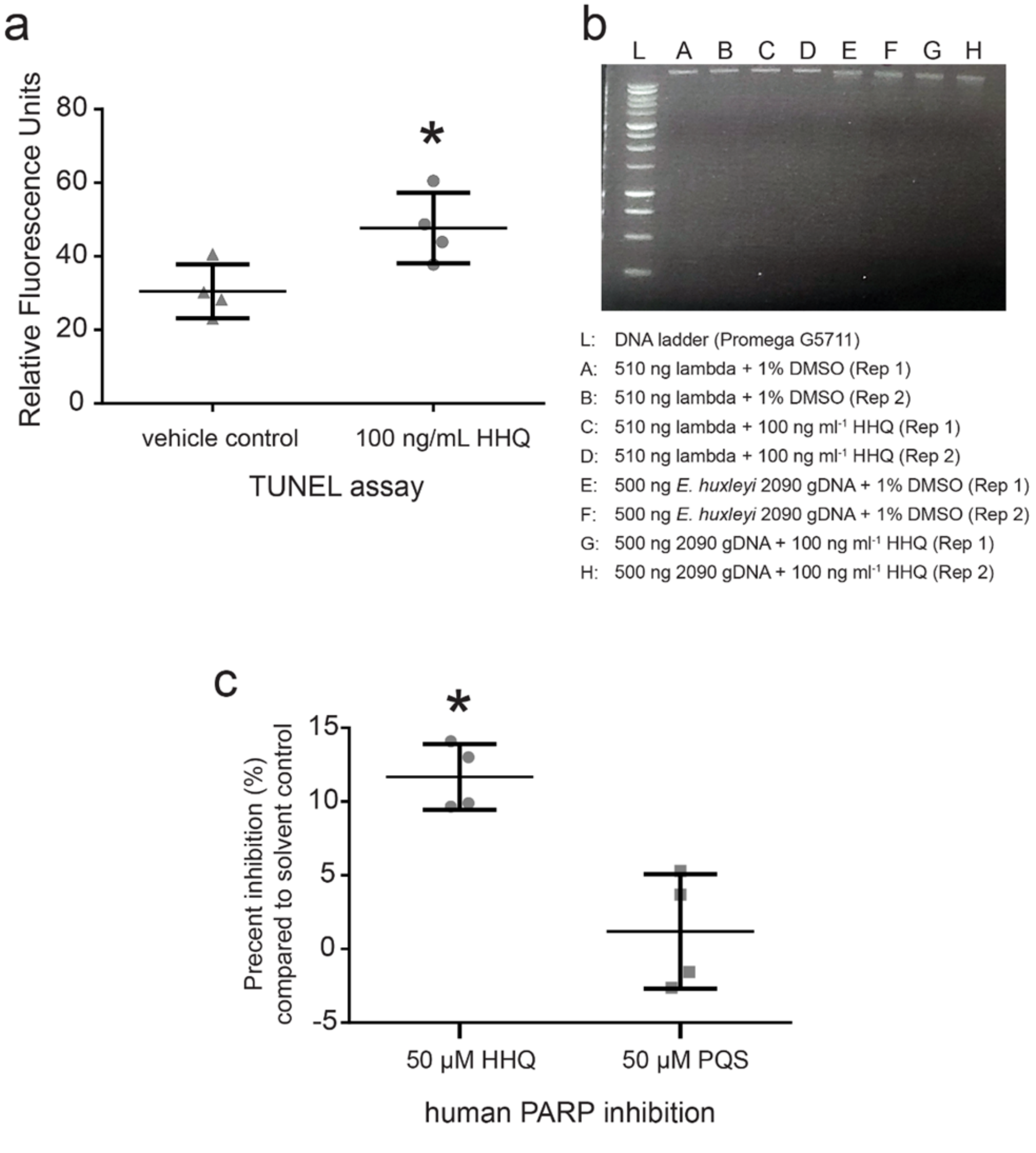

**Figure S16.** Protein homology modeling of *E. huxleyi* PARP1-like homolog (XP_005783504.1) to the crystal structure of human PARP1 (PDB 2RD6) with the PARP inhibitor veliparib. (a) The sequence of *E. huxleyi* PARP-like homolog (XP_005783504.1) was aligned to human PARP1 (PDB 2RD6). Two key amino acid residues (red), the Tyr and Glu, are strictly conserved between human PARP and *E. huxleyi* protein. (b, c) The active site of PDB 2RD6 shows the binding of the inhibitor veliparib. Regions in 2RD6 with greatest homology to *E. huxleyi* PARP-like homolog are colored in pink with the remainder of the protein chain in green. The region comprising the active site for small molecule binding is highly conserved between the two proteins with two key binding interactions observed – the near co-planar arrangement of a tyrosine side chain phenyl to the inhibitor, and a water mediated hydrogen bond of the basic nitrogen atom of the inhibitor to a glutamate side chain carboxyl (c).

a

```
Query  750   SSLPPPVARFVSLIGDKKMMAAQLQEMHVDLEKMPLGSISTAQIKRGYEALTRVATQLQT  809
             S LP PV   + +I D + M   + E  +DL+KMPLG +S  QI+  Y  L+ V   +
Sbjct  2     SKLPKPVQDLIKMIFDVESMKKAMVEYEIDLQKMPLGKLSKRQIQAAYSILSEVQQAVSQ  61

Query  810   PAADGDEQKHRLLTLTTHFYTTIPHKFGLDEVPPVIDSLPMLKDKLDMVEALLQVQESQK  869
              ++D      ++L L+  FYT IPH FG+ + PP++++   ++ K +M++ LL ++ +
Sbjct  62    GSSDS-----QILDLSNRFYTLIPHDFGMKK-PPLLNNADSVQAKAEMLDNLLDIEVAYS  115

Query  870   LSKSLGPAAEQHPLDAMYAQLQCGL--APASEAEHEMVRRYAANTHGSTHDTYKINVKDV  927
             L +     + + P+D  Y +L+   +        E E++R+Y  NTH +TH+ Y + V D+
Sbjct  116   LLRGGSDDSSKDPIDVNYEKLKTDIKVVDRDSEEAEIIRKYVKNTHATTHNAYDLEVIDI  175

Query  928   LAVSRAAEQPGFK--ASLANRQLLWHGSRLTNWAGILSQGLRIAPPEAPVTGYMFGKGVY  985
               + R  E   +K    L NR+LLWHGSR TN+AGILSQGLRIAPPEAPVTGYMFGKG+Y
Sbjct  176   FKIEREGECQRYKPFKQLHNRRLLWHGSRTTNFAGILSQGLRIAPPEAPVTGYMFGKGIY  235

Query  986   FADMSSKSANYCGATSANPTAVLLLCDVALGSQYERLNAEYEAAESCKKAGKASTWGVGK  1045
             FADM SKSANYC  +  +P  ++LL +VALG+ YE  +A +    S    GK S  G+GK
Sbjct  236   FADMVSKSANYCHTSQGDPIGLILLGEVALGNMYELKHASH---ISKLPKGKHSVKGLGK  292

Query  1046  FAPADGGASSIEGGAIRVPMGEAGDSEYMKANFARLQKAEGGKARAALLYNEYIVYDVSQ  1105
               P      S++G  + VP+G    S               G    +LLYNEYIVYD++Q
Sbjct  293   TTPDPSANISLDG--VDVPLGTGISS---------------GVNDTSLLYNEYIVYDIAQ  335

Query  1106  IQMKYVVVCELEF  1118
             + +KY++  +  F
Sbjct  336   VNLKYLLKLKFNF  348
```

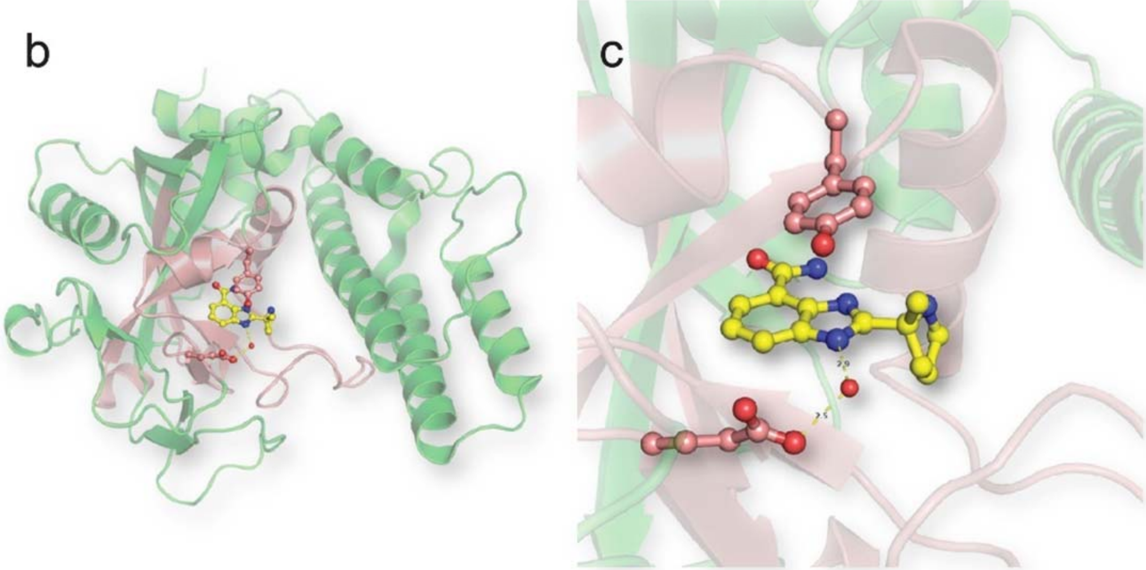

**Figure S17.** The Ocean Gene Atlas was mined for homologs of the HHQ biosynthesis and transcriptional regulation genes (Genbank KT879191 thru KT879199; corresponding to (a) thru (h)) matching the query from *Pseudoalteromonas piscicida* (A757) (Harvey et al. 2016). Circle diameter indicates the percent of genes in the sample and circle color denotes initial sampling fraction. Specific e-value cut-offs for each gene are indicated in each figure.

E. L. Harvey et al., A Bacterial Quorum-Sensing Precursor Induces Mortality in the Marine Coccolithophore, *Emiliania huxleyi*. *Front Microbiol* 7 (2016). doi: 10.3389/fmicb.2016.00059

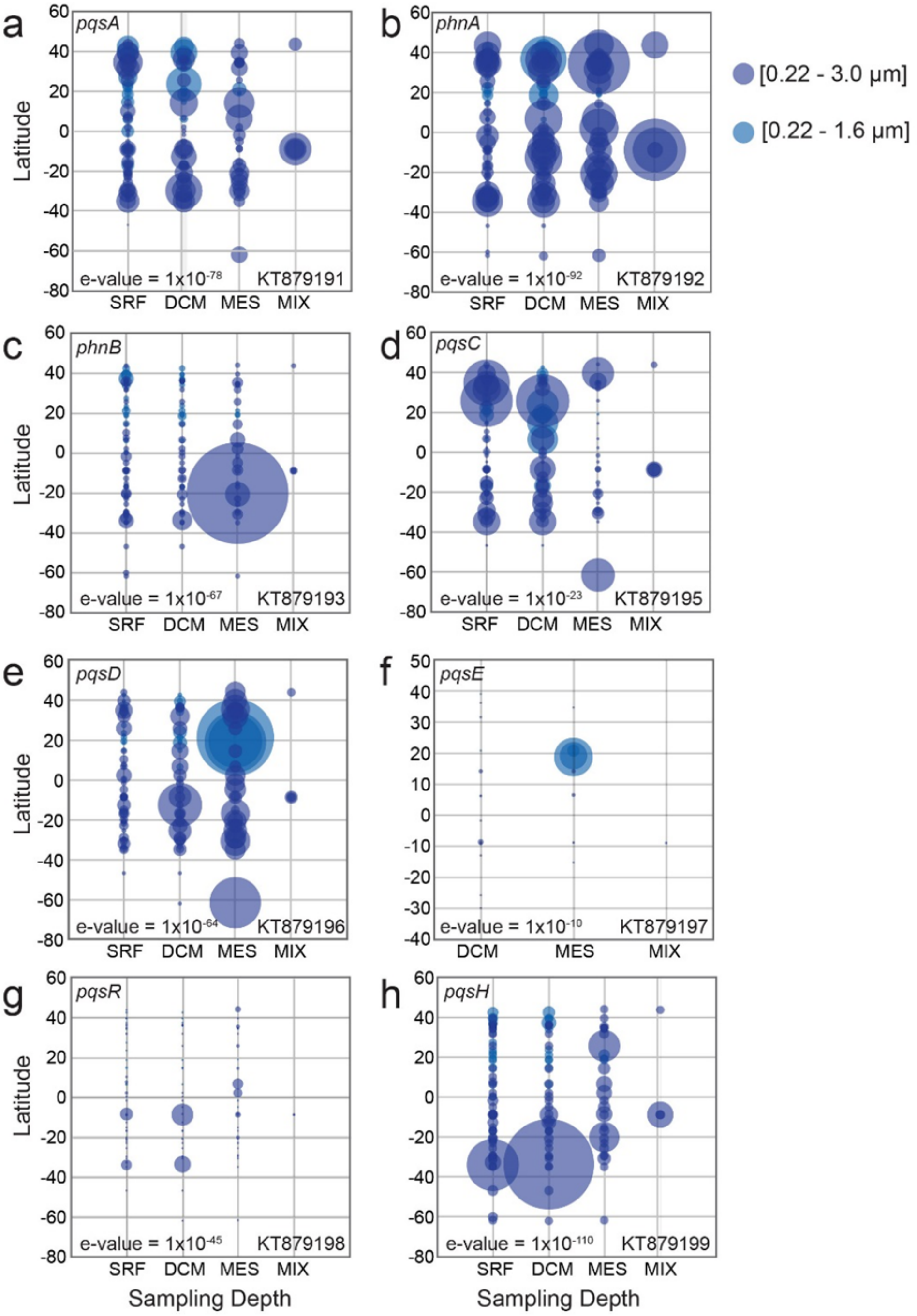

**Figure S18.** Composite figure showing the detection of HHQ in environmental samples. The extracted ion chromatograms for HHQ ($[M+H]^+$ = 244.17 *m/z*) detected by high-resolution (450K) orbitrap LC-ESIMS for the standard (a) and environmental samples (b). HHQ confirmation via MS/MS analysis in environmental samples compared to an authentic standard. High mass accuracy HCD fragmentation spectra (35% collision energy) of the $[M+H]^+$ ion were collected at 38 minutes (c).

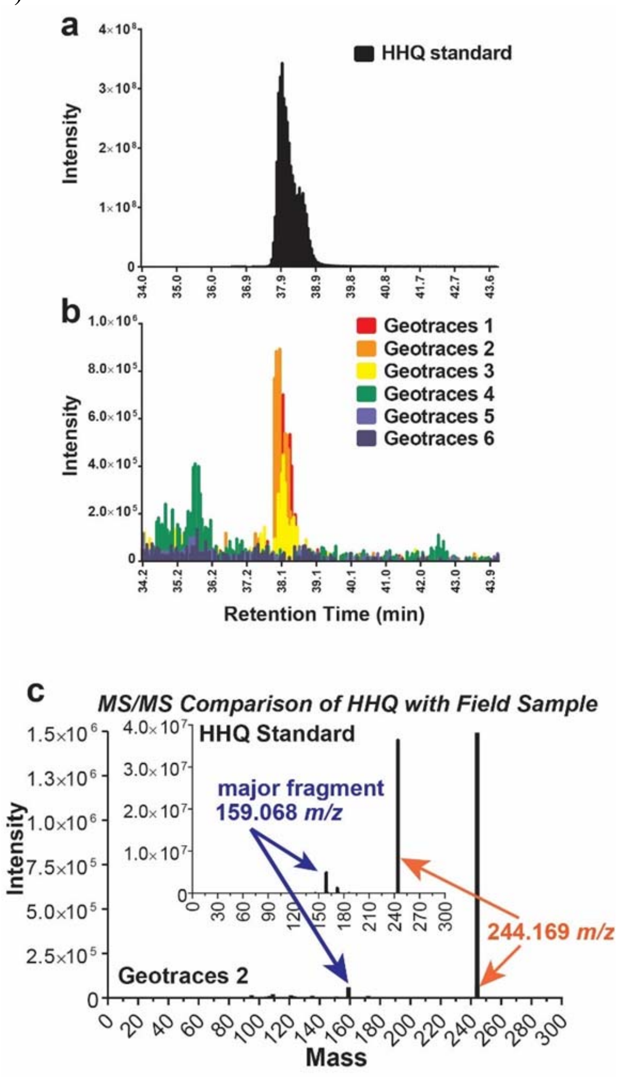

**Table S1.** Quantification of differentially expressed transcripts and proteins following HHQ exposure for 24 and 72 h.

For padj <0.05

| Transcript | 24 hr | | | 72 hr | | |
|---|---|---|---|---|---|---|
| | Up | Down | Unchanged | Up | Down | Unchanged |
| Low | 13 | 20 | 31476 | 0 | 0 | 31549 |
| Medium | 2,702 | 1,990 | 26817 | 382 | 159 | 31008 |
| High | 5,948 | 6,605 | 18956 | 6,166 | 5,698 | 19685 |

| Protein | 72 hr | | |
|---|---|---|---|
| | Up | Down | Unchanged |
| Low | 0 | 0 | 5528 |
| Medium | 3 | 0 | 5525 |
| High | 628 | 375 | 4525 |

**Table S2.** Purity broth recipe.

| **f/2MB purity broth** | **f/2MM purity broth** |
|---|---|
| 75% sterile seawater base (25% ultrapure water) with: | 100% sterile seawater base with: |
| $NH_4Cl$…800 µM | $NH_4Cl$…800 µM |
| $NaNO_3$…882 µM | $NaNO_3$…882 µM |
| $NaH_2PO_4$…36.2 µM | $NaH_2PO_4$…36.2 µM |
| Marine Broth 2216…1.7 g/L | Organics: |
| | pyruvate , acetate, glycerol …0.05% (w/v) |
| *Trace metals:* | *Trace metals:* |
| $FeCl_3$ $6H_2O$ …11.7 µM | $FeCl_3$ $6H_2O$ …11.7 µM |
| $Na_2EDTA$ $2H_2O$ …11.7 µM | $Na_2EDTA$ $2H_2O$ …11.7 µM |
| $CuSO_4$ $5H_2O$ …39.3 nM | $CuSO_4$ $5H_2O$ …39.3 nM |
| $Na_2MoO_4$ $2H_2O$ …26.0 nM | $Na_2MoO_4$ $2H_2O$ …26.0 nM |
| $ZnSO_4$ $7H_2O$ …76.5 nM | $ZnSO_4$ $7H_2O$ …76.5 nM |
| $CoCl_2$ $6H_2O$ …42.0 nM | $CoCl_2$ $6H_2O$ …42.0 nM |
| $MnCl_2$ $4H_2O$ … 0.91 µM | $MnCl_2$ $4H_2O$ …0.91 µM |
| *Vitamins:* | *Vitamins:* |
| Thiamine HCl (vit. $B_1$) …296 nM | Thiamine HCl (vit. $B_1$) …296 nM |
| Biotin (vit. H) …2.05 nM | Biotin (vit. H) …2.05 nM |
| Cyanocobalamin (vit. $B_{12}$) …0.369 nM | Cyanocobalamin (vit. $B_{12}$) …0.369 nM |

**Dataset S1.** Master list of all protein coding genes identified in the transcript and proteomic analysis. Relative abundance values for all transcripts detected in this study. Columns 35 through 71 represent the regularized log transformation (rlog) normalized counts for each transcript. NA indicates this transcript was not detected. Columns 72 through 97 contain mean normalized counts used for dispersion estimation and information related to fold change differences derived from DESeq2 to determine statistically significantly different transcript abundances. Abundance values for all protein detected. Columns 98 through 115 represent the peak area for each protein. NA indicates this protein was not detected. Columns 116 through 128 contain average protein abundance and information related to fold change differences between protein abundances. Column 130 contains the potential identity of the gene/protein determined via homology using the NCBI protein BLAST tool (e-value < $10^{-20}$).

**Dataset S2.** Lists of gene ontology (GO) IDs used to categorize uncharacterized *E. huxleyi* proteins. An initial GO ID was selected and then all GO IDs below that term in the GO hierarchy are listed.
